# Mice lacking *Astn2* have ASD-like behaviors and altered cerebellar circuit properties

**DOI:** 10.1101/2024.02.18.580354

**Authors:** Michalina Hanzel, Kayla Fernando, Susan E. Maloney, Shiaoching Gong, Kärt Mätlik, Jiajia Zhao, H. Amalia Pasolli, Søren Heissel, Joseph D. Dougherty, Court Hull, Mary E. Hatten

**Affiliations:** Laboratory of Developmental Neurobiology, The Rockefeller University, New York, NY, USA 10065; Neurobiology Department, Duke University, Durham, NC, USA; Dept of Psychiatry and the Intellectual and Developmental Disabilities Research Center, Washington University Medical School, St Louis, MO, USA; Dept of Genetics, Washington University Medical School, St Louis, MO, USA; Weill Cornell Medical College, New York, NY, USA; Electron Microscopy Resource Center, The Rockefeller University, New York, NY, USA 10065; Proteomics Resource Center, The Rockefeller University, New York, NY, USA 10065

## Abstract

Astrotactin 2 (ASTN2) is a transmembrane neuronal protein highly expressed in the cerebellum that functions in receptor trafficking and modulates cerebellar Purkinje cell (PC) synaptic activity. We recently reported a family with a paternally inherited intragenic *ASTN2* duplication with a range of neurodevelopmental disorders, including autism spectrum disorder (ASD), learning difficulties, and speech and language delay. To provide a genetic model for the role of the cerebellum in ASD-related behaviors and study the role of ASTN2 in cerebellar circuit function, we generated global and PC-specific conditional *Astn2* knockout (KO and cKO, respectively) mouse lines. *Astn2* KO mice exhibit strong ASD-related behavioral phenotypes, including a marked decrease in separation-induced pup ultrasonic vocalization calls, hyperactivity and repetitive behaviors, altered social behaviors, and impaired cerebellar-dependent eyeblink conditioning. Hyperactivity and repetitive behaviors were also prominent in *Astn2* cKO animals. By Golgi staining, *Astn2* KO PCs have region-specific changes in dendritic spine density and filopodia numbers. Proteomic analysis of *Astn2* KO cerebellum reveals a marked upregulation of ASTN2 family member, ASTN1, a neuron-glial adhesion protein. Immunohistochemistry and electron microscopy demonstrates a significant increase in Bergmann glia volume in the molecular layer of *Astn2* KO animals. Electrophysiological experiments indicate a reduced frequency of spontaneous excitatory postsynaptic currents (EPSCs), as well as increased amplitudes of both spontaneous EPSCs and inhibitory postsynaptic currents (IPSCs) in the *Astn2* KO animals, suggesting that pre- and postsynaptic components of synaptic transmission are altered. Thus, ASTN2 regulates ASD-like behaviors and cerebellar circuit properties.

## INTRODUCTION

ASTN2 is a vertebrate-specific neuronal glycoprotein with important roles in trafficking proteins including synaptic receptors ^1^, as well as the neuron-glial adhesion protein ASTN1 that functions in glial-guided neuronal migration ^2–5^. ASTN2 expression levels are highest in cerebellar Purkinje cells (PCs) and granule neurons, with lower levels of expression in the cortex, olfactory bulb, and dentate gyrus of the hippocampus ^1,4^. Copy number variations (CNVs) in *ASTN2* have been identified as a significant risk factor for ASD^6–8^, and *ASTN2* is listed as a gene implicated in ASD susceptibility by the SFARI initiative of the Simons Foundation. Importantly, we recently reported a family with a paternally inherited intragenic *ASTN2* duplication, which results in *ASTN2* haploinsufficiency. The family manifests a range of neurodevelopmental disorders, including ASD, learning difficulties, and speech and language delay^1^. The high levels of ASTN2 expression in the mouse cerebellum suggest that *ASTN2* mutations such as those found in patients with *ASTN2* CNVs could lead to altered cerebellar function. Furthermore, our cellular and molecular studies on mouse cerebellum show that ASTN2 binds to and regulates the trafficking of multiple synaptic proteins, many of which are implicated in ASD, and modulates cerebellar PC synaptic activity^1^. We therefore generated novel *Astn2* KO and cKO mouse lines to examine whether ASTN2 loss affects ASD-associated behaviors and cerebellar circuit properties.

Although the cerebellum has long been considered to be a purely motor structure, recent studies reveal that it also has critical nonmotor functions, including language^9,10^, social cognition^11^, and emotional processing^12,13^. Cerebellar activation is observed in humans during social cognition tasks^14^, and stimulation of the mouse cerebellum modulates dopamine release in the medial prefrontal cortex of wild-type animals^15^, but not in several mouse models of ASD^16^. Recent studies demonstrate a direct, monosynaptic pathway from the cerebellum to the ventral tegmental area that controls social behaviors^17^ and show that the cerebellar-prefrontal cortex circuits mediate cerebellum-regulated social and repetitive/inflexible behaviors^18^. Moreover, a loss of cerebellar PCs is one of the most consistent structural findings in postmortem studies of patients with ASD^19^ and cerebellar injury at birth results in an approximately 40-fold increase in ASD by age 2^20,21^. In addition, specific targeting of cerebellar PCs in mouse models of ASD-associated genes leads to impaired cerebellar learning^22^ and social behaviors^23–25^. Thus, cerebellar dysfunction is strongly implicated in ASD.

In the present study, we generated global and conditional loss of function *Astn2* mouse lines and analyzed ASD-related behaviors and cerebellar circuit properties. We found that mice lacking *Astn2* have deficits in social behaviors and in the number and properties of ultrasonic vocalization (USV) calls, as well as an increase in repetitive behaviors and hyperactivity, all behavioral changes that are characteristic of ASD^26^. Importantly, the hyperactivity and repetitive behavior changes were also found in a mouse line with a conditional loss of *Astn2* specific to PCs, suggesting a role for the cerebellum in these behaviors. Consistent with this finding, the ASD-like behaviors were accompanied by changes in the structure of PC dendritic spines in the posterior vermis and Crus1 of *Astn2* KO animals, areas associated with repetitive behaviors and social behaviors, respectively^18^. Molecular studies revealed an upregulation of the neuron-glial adhesion protein ASTN1 with concomitant changes in the volume of Bergmann glial fibers in the molecular layer. Finally, we also observed changes in cerebellar circuit properties evidenced by changes in spontaneous EPSCs and IPSCs. These studies suggest that ASTN2 functions in ASD-like behaviors and cerebellar circuit properties, and show, in agreement with other studies on cerebellar circuits in ASD^24^, that even subtle changes in cerebellar anatomy and physiology can lead to significant behavioral changes.

## RESULTS

### Generation of *Astn2* global and conditional knockout lines

The *Astn2* knockout (KO) mouse line was generated using CRISPR-Cas9 technology by targeting the first exon and the promoter region of the *Astn2* gene (see the Methods section for details). The loss of ASTN2 protein expression was confirmed using Western blot analysis (Supp. Fig. 1) and the proteomic analysis of whole cerebellar tissue (Fig. 5). *Astn2* KO mice did not show reduced survival or fertility and their weights were comparable across genotypes at P22. The gross morphology of the cerebellum, including the size and weight, foliation pattern, cell densities and layer formation, was normal (Supp. Fig. 2). We did not observe any abnormalities in cerebellar neurogenesis, granule cell proliferation or glial-guided neuronal migration (Supp. Fig. 3). Next, we generated a conditional *Astn2* knockout mouse line by inserting two loxP sites flanking exon 1 and the promoter region of *Astn2* gene using a similar approach as with the global KO mouse (see Methods). The *Astn2^flox/flox^* line was then crossed with the *Pcp2*-*Cre* line (Gensat) to generate a PC-specific conditional knockout line (*Astn2* cKO, Supp. Fig. 1). *Astn2* cKO mice did not show reduced survival or fertility, their weights were comparable across genotypes at P22 and no gross abnormalities were observed.

### *Astn2* KO pups produce fewer and less complex ultrasonic vocalizations in a maternal separation assay

Individuals with *ASTN2* CNVs have language deficits^1^, and many mouse models of ASD show deficits in USVs in social communication^27–29^. To understand the influence of *Astn2* loss of function on early social communication, we measured pup isolation calls induced by maternal isolation (Fig. 1A). USVs were recorded after pup isolation at postnatal days (P)5 – P14. Loss of *Astn2* expression robustly influenced USV production (Fig. 1B). Specifically, *Astn2* KO mice exhibited significantly fewer USVs during the 5-minute recording session across P6-10 (Fig. 1C). In addition, heterozygous mutants (Het) also showed a significant reduction in call number compared to wild types on P6 – P8, demonstrating an intermediate phenotype. This phenotype did not interact with sex (data not shown). We next examined the temporal and spectral features of the calls to determine if the calls varied following the loss of *Astn2*. Complementary to the number of individual calls, the *Astn2* KO mice also produced fewer bouts of calls, defined as a sequence of calls separated by pauses <0.5 seconds, compared to Het and WT littermates on days P6-10. Again, Hets showed an intermediate phenotype with reduced bouts only on day 8 (Fig. 1D). In addition, *Astn2* KO mice exhibited longer pauses between calls within a bout compared to WT & Het littermates overall, with a particularly robust decrease at P10 (Fig. 1E). We found that overall *Astn2* KO mice produced shorter individual calls compared to both heterozygous *Astn2*^+/-^ and WT littermates (Fig. 1F). We also examined spectral features of the calls. We observed a significant narrowing of frequency pitch range in all the calls produced by *Astn2* KO mice at ages P6-10 compared to those produced by their heterozygous *Astn2* and WT littermates (Fig. 1G). No differences were observed for the average frequency pitch or the peak amplitude of the sound pressure level (loudness) of all calls produced by the mice (data not shown). Next, we separated the calls into those that contain a frequency pitch jump, known as dynamic calls, and those that do not contain a pitch jump, known as flat calls. We observed a significant decrease in the fraction of dynamic calls produced by *Astn2* KO mice, which was particularly robust P6 – P10, compared to heterozygous *Astn2* and WT littermates (Fig. 1H). The average pitch and pitch range of dynamic calls were not different between groups (data not shown). While the flat calls produced by *Astn2* KO did not differ in their average pitch, they were significantly narrowed in their pitch range compared to heterozygous *Astn2* and WT littermates, particularly at P7-10 (Fig. 1I). In conclusion, a loss of *Astn2* in the mouse results in a robust decrease in USV production, particularly during the peak age range for this behavior (P6-P10), and significantly influences the spectral pitch range of the produced calls.

**Figure 1:**
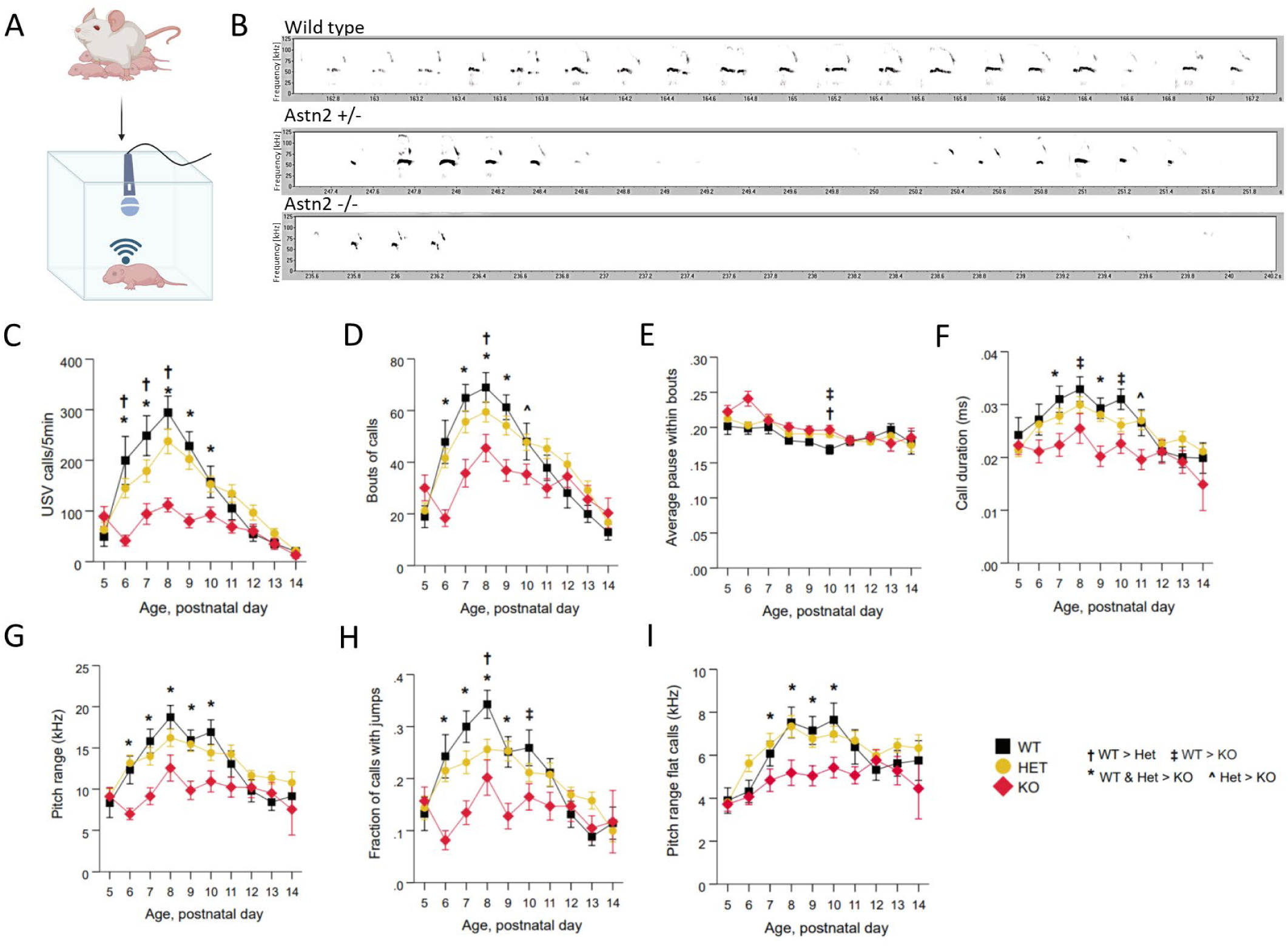
Loss of *Astn2* results in fewer and less dynamic ultrasonic vocalizations in the mouse pups. **A)** Pups were isolated from their mothers at postnatal days (P)5 – P14 and placed in a soundproof chamber fitted with an ultrasonic microphone. Ultrasonic vocalizations (USVs) were recorded for 5 mins. **B)** Sample spectrograms illustrating the differences in USVs from P7 wild type (WT) (n = 20), heterozygous (HET) (n = 52) and *Astn2* KO (n = 24) animals (frequency in kHz as a function of time). Loss of ASTN2 expression robustly influenced USV production and dynamics on a number of measures at P6-P10: **C)** Number of USV calls during the 5-minute recording session was reduced in *Astn2* KOs. Heterozygous animals showed a significant reduction at P6-P8. **D)** Bouts of calls were reduced in *Astn2* KOs. **E)** Average pause within bouts was reduced in *Astn2* KOs at P10. **F)** Call duration was reduced in *Astn2* KOs. **G)** Pitch range of all USV calls was reduced in *Astn2* KOs. **H)** Fraction of calls that are dynamic (contain a pitch jump) was reduced in *Astn2* KOs. Heterozygous animals show a reduction in dynamic calls at P8. **I)** Range of frequency pitch for flat calls was reduced in *Astn2* KOs at P7-P10.

### *Astn2* KO animals show defects in social behaviors in a three-chamber test

Individuals with ASD show abnormalities in social behavior. To understand the influence of *Astn2* on social behaviors, we evaluated social behaviors in adult mice (8-12 weeks of age) in the three-chamber sociability and social novelty tests^30^. In this paradigm, the test mouse is placed in the middle chamber of a three-chamber apparatus with two empty wire baskets in the other two chambers (Fig. 2A). The test consists of three phases starting with an initial 10-minute habituation period to the empty baskets. In the second phase, a social stimulus (a novel mouse) is placed in one of the baskets and a non-social stimulus, such as a Lego block, is placed in the other basket. The time that the test mouse spends investigating the two stimuli is measured. A typical wild type mouse prefers to spend time with a social stimulus. In the third phase, the familiar mouse is left in their basket and the non-social stimulus is exchanged for a novel mouse. In this case, a typical wild type mouse prefers to spend time with a novel mouse instead of the familiar mouse. In our experiments, *Astn2* KO mice showed an overall preference for a social, rather than a non-social stimulus, similar to their wild type littermates. However, they spent significantly less total time investigating the social stimulus compared to wild type mice (Fig. 2B). Additionally, while wild type littermates spent significantly longer investigating the novel mouse, *Astn2* KOs did not show a preference for the novel mouse and spent the same amount of time investigating the novel and the familiar counterparts (Fig. 2C). Therefore, *Astn2* KO mice exhibit abnormal social behavior.

**Figure 2:**
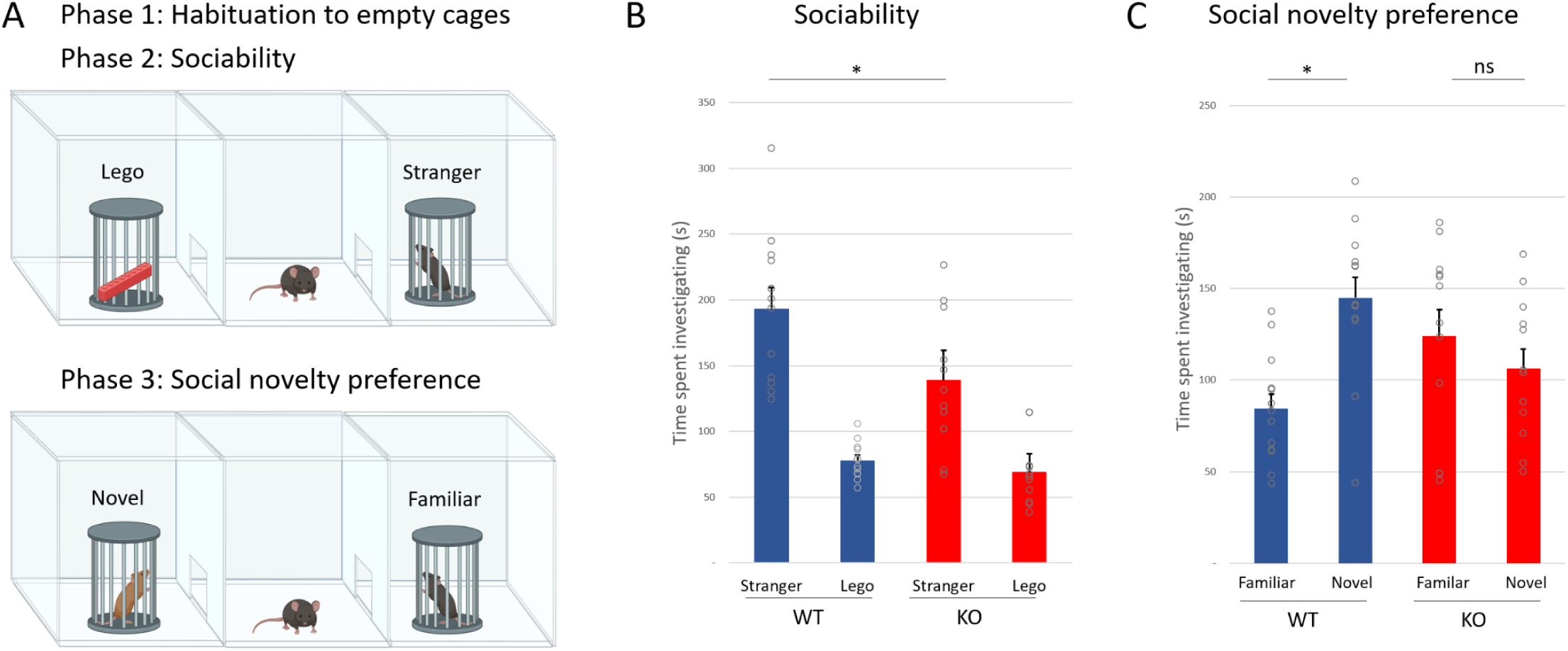
Social behavior is altered in the *Astn2* KO mice. **A)** The three-chamber social test was performed on 8–12-week-old animals (n = 12 WT, 11 KO). The test consisted of three phases, starting with a 10-minute habituation to the empty cages, followed by two 10-minute testing phases. In phase 2, to test sociability, mice were exposed to a non-social (Lego block) and a social stimulus (stranger mouse). In phase 3, to test social novelty preference, the non-social stimulus was replaced by a novel mouse. The familiar mouse from phase 2 was left in their cage. The time spent interacting with the stimuli was manually scored by the investigator. **B)** In the first phase of the experiment, WT mice show a strong preference for the social stimulus (p <0.0001). *Astn2* KO animals also show a preference for the social stimulus (p= 0.001), however, the absolute time they spend interacting with the social stimulus is significantly reduced as compared to the WT (p= 0.027). **C)** In the third phase of the experiment, WT animals interact with the novel animals significantly longer than with the familiar animals (p= 0.0003). In contrast, the *Astn2* KO animals do not show a preference for the novel animal and spend as much time with the familiar animal as with the novel animal (p = 0.35). Data is presented as the mean + SEM. *p < 0.05, **p < 0.01, ***p <0.001, ****p <0.0001, ns= not significant

### *Astn2* KO animals are hyperactive and show repetitive behaviors and reduced anxiety in an open field assay

One of the most prominent features of individuals with ASD is repetitive behavior. Some patients also show hyperactivity and/or an increase in anxiety. We used the open field experimental assay to measure these behaviors in *Astn2* KO mice. 8–12-week-old animals were placed in an open arena for 1 hour and allowed to explore freely (Fig. 3A). We found that *Astn2* KO mice travel a significantly longer distance in the arena compared to their wild type and heterozygous littermates (Fig. 3B). Additionally, *Astn2* KO mice display a number of behaviors that are suggestive of repetitive behaviors, including increased rearing (standing vertically on their hind paws) (Fig. 3C) and an increase in the number of revolutions (circling in place) (Fig. 3D). Anxiety-like phenotypes can also be measured in the open field paradigm. Typically, mice prefer to spend more time toward the periphery of the arena and avoid the center. Spending a higher proportion of time in the periphery is indicative of heightened anxiety-like avoidance behaviors and spending more time in the center is indicative of lower anxiety. *Astn2* KO animals spent more time in the center of the arena compared to WT littermates (Fig. 3E) suggesting lowered anxiety. An additional measure of anxiety-like behavior involves using the light/dark box where half of the arena is covered by a dark box (Fig. 3F). Mice that prefer to spend more time in the dark versus the light compartment or take longer to enter the light area are exhibiting more avoidance behavior. *Astn2* KO mice did not show a preference for the dark compartment (Fig. 3G) but had lower latency to enter the light compartment compared to control littermates (Fig. 3H). Therefore, *Astn2* KO mice exhibit hyperactivity and repetitive behaviors and display lowered anxiety phenotypes.

**Figure 3:**
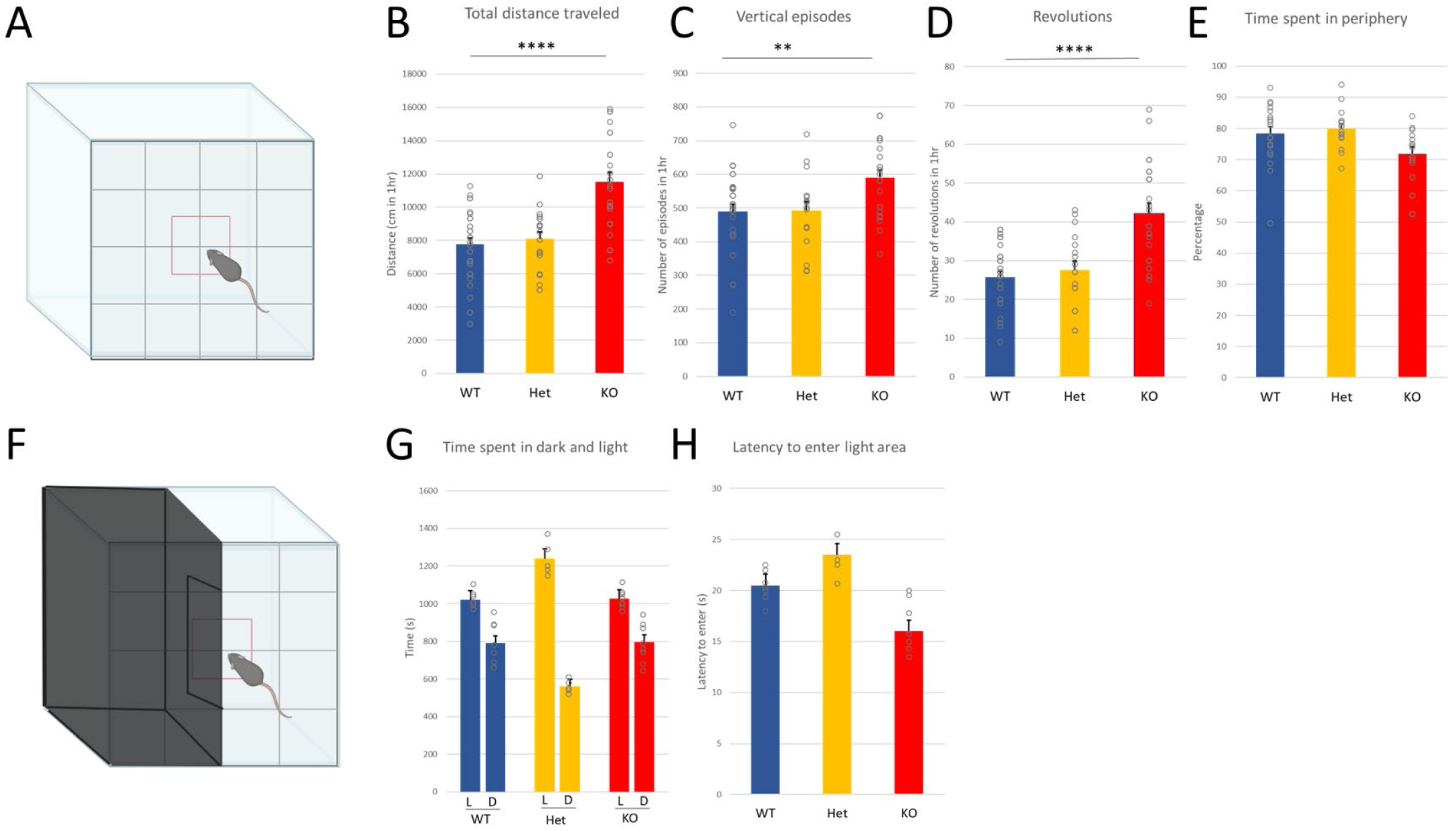
*Astn2* KO mice show hyperactivity and repetitive behaviors, but not anxiety, in the open field test. **A)** 8-12 weeks old wild type (n = 29), heterozygous (n = 17) and *Astn2* KO (n = 23) animals were placed in an open field arena for 1 hour and allowed to explore freely. The center of the arena was assigned in the software. **B)** The total distance traveled was increased significantly in the KO animals (p < 0.0001). **C)** The number of vertical episodes (rearing) was significantly increased in the *Astn2* KOs as compared to WTs and Hets (p = 0.005). **D)** The number of revolutions (circling) was significantly increased in the *Astn2* KOs (p < 0.0001). **E)** Time spent in the center versus the periphery of the arena was measured. *Astn2* KOs spend less time in the periphery of the arena (p = 0.026). **F)** A subset of animals (n = 7 WT, 5 Het, 8 KO) were tested in a light/dark open field paradigm where a black box covers half of the arena. **G)** All genotypes spent significantly more time in the light part of the arena (L) versus the dark part (D) and there were no differences in the amount to time spent in the light (L) versus the dark (D) between WT and KO animals (p = 0.9). Het animals spent significantly more time in the light compared to WT and KO animals (p < 0.0001). **H)** The latency to enter the light compartment significantly decreased in the *Astn2* KO (p < 0.0001). Data is presented as the mean + SEM. Data analyzed with One way ANOVA with Tukey Kramer post-hoc test *p < 0.05, **p < 0.01, ***p <0.001, ****p <0.0001

### *Astn2* KO animals show abnormal cerebellar-dependent associative learning

Individuals with ASD often show abnormal responses to eyeblink conditioning, an associative learning paradigm in which cerebellar circuitry plays a central role^31–34^. To assess cerebellar-dependent associative learning, we trained adult (8-15 weeks old) *Astn2* KO mice and age-matched, WT littermate controls on a delay eyeblink conditioning task^35^. Mice were head-fixed on a freely moving cylindrical treadmill (Fig. 4A) and conditioned with a 250-ms light stimulus (conditioned stimulus (CS), blue light) and a co-terminating, 30-ms corneal air puff (unconditioned stimulus (US), 30 PSI) (Fig. 4 A,B). Across trials, conditioned responses (CRs) developed as a predictive eyelid closure preceding the US (Fig. 4 B). On average, we observed a trend toward reduced learning in *Astn2* KO animals relative to controls, with a decrease in CR probability (Fig. 4 C, *top*) and amplitude (Fig. 4 C, *middle*) at later learning timepoints. Notably, however, these averages obscure the variability across individual animals. While all WT animals exhibited learning, 5 of 9 *Astn2* KO animals exhibited little or no learning (Fig. 4 C, *bottom*, 4 D, *bottom*). In addition, the *Astn2* KO animals that did learn exhibited altered eyeblink kinematics, displaying more average peaks in their conditioned responses (Fig. 4 D). These data are consistent with individuals that have ASD, who often produce abnormal conditioned responses after delay eyeblink conditioning, frequently resulting in eyelid closures that have altered kinematics^33^.

**Figure 4.**
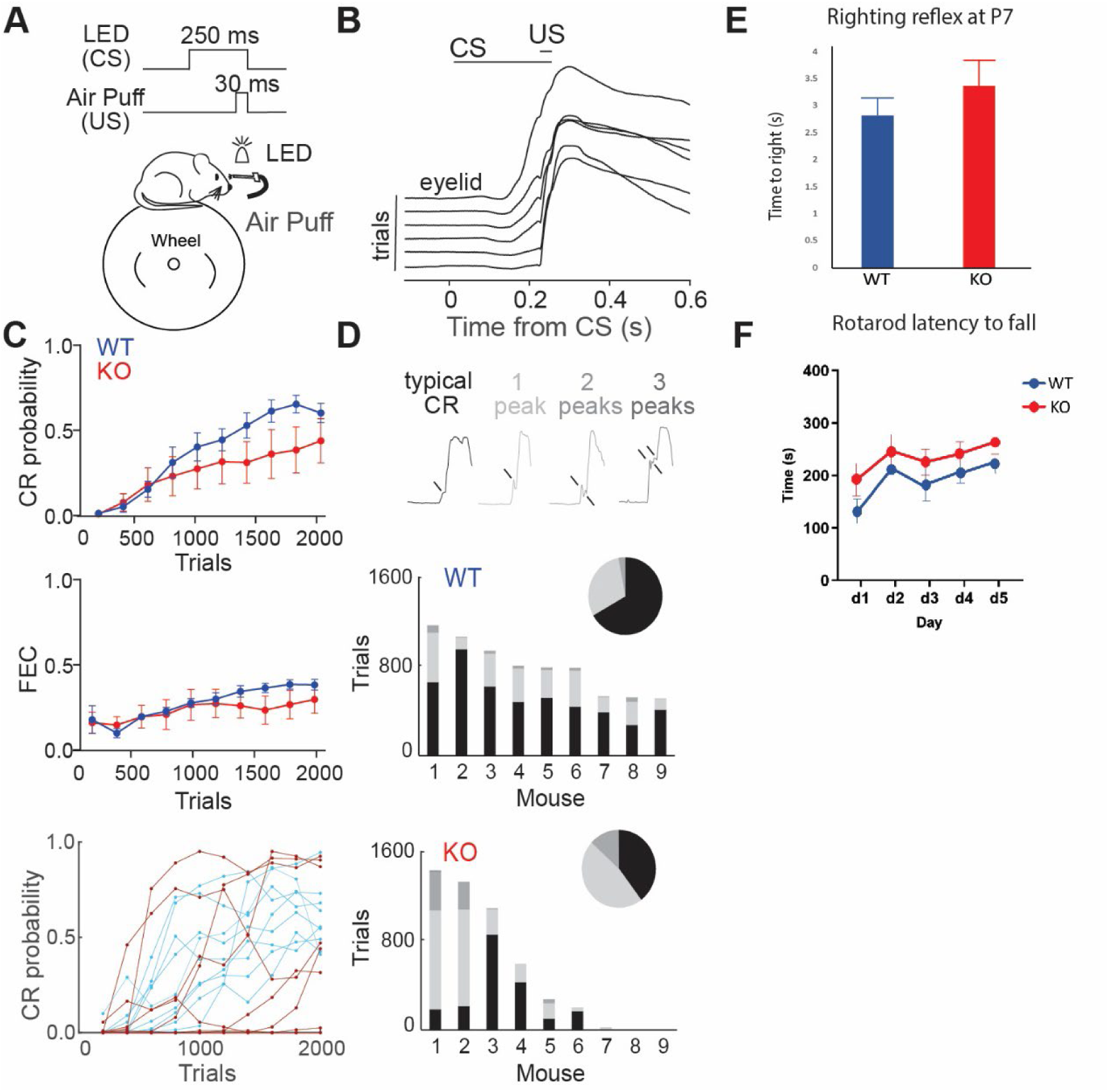
*Astn2* KO mice show abnormal cerebellar-dependent associative learning but normal righting reflex and rotarod behavior. **A)** Behavioral configuration for delay eyeblink conditioning. Mice (8-15 weeks old, N = 9/genotype) were head-fixed and could run freely on a cylindrical treadmill while high-speed videography recorded eyelid position during stimuli presentation. **B)** Selected average eyelid traces (100-trial average) over the course of training from an example WT mouse in response to a 250-ms LED conditioned stimulus (CS) co-terminating with the delivery of a 30-ms aversive air puff unconditioned stimulus (US). **C)** *Top,* binned conditioned response (CR) probability over training (200-trial average). *Middle,* binned CR amplitude (fraction of eyelid closed (FEC), 200-trial average) over training with CRs preserved in trial space. *Bottom,* CR probability plotted for individual WT (light blue) and KO (maroon) mice. **D)** *Top,* example eyelid traces showing various CR topographies observed during CS-US trials. CRs with three peaks comprised <1% of all observed CRs. *Middle insert,* detected CR peaks from all CS-US trials of all WT mice (N = 7128 trials). *Middle,* proportional contribution of individual WT mice. *Bottom inset,* same as *middle inset,* for all CS-US trials of all KO mice (N = 4943 trials). *Bottom,* same as *middle,* for individual KO mice. **E)** P7 wild type (n = 12) and Astn2 KO (n = 13) pups were evaluated using the righting reflex assay by placing pups in a supine position. Time to completely right themselves was recorded. No difference was found between WT and KO animals. **F)** 8-12 week old wild type (n = 8) and Astn2 KO (n = 10) animals were placed on an accelerating rotarod for five consecutive days (d). An average of three trials per day was recorded. Time to fall from the rotarod was measured. WT and *Astn2* KO animals did not show significantly different results in their latency to fall. Data is presented as the mean ± SEM.

### *Astn2* KO animals show normal righting reflex and rotarod behavior

As mutations in cerebellar genes often result in motor abnormalities in mice, we investigated motor behavior by testing pup righting reflex and adult mice performance on the accelerating rotarod test. P7 pups were placed on their backs in a supine position. The time taken for the mice to right themselves was measured. *Astn2* KO pups did not significantly differ compared to control mice in the time taken to right themselves (Fig. 4 E). Next, adult 8–12-week-old mice were placed on a rotarod that accelerated from 4 to 40 RPM in 5 min and were tested on five consecutive days with three trials each day. *Astn2* KO animals did not show defects in balance and coordination as their performance did not significantly differ compared to WT littermates (Fig. 4F). Thus, while *Astn2* KO animals have ASD-like behaviors they do not exhibit righting reflex deficits or deficits in motor coordination and balance on the rotarod.

### Purkinje cell-specific deletion of *Astn2* recapitulates some ASD-like behaviors

The *Astn2^flox/flox^* line was crossed with the *Pcp2*-*Cre* line to generate a PC-specific conditional knockout line. We used this line to study the effect of the loss of *Astn2* in PCs on activity, repetitive behaviors, social behavior, and motor behavior. We found that *Astn2* cKO animals recapitulated behavior seen in the global *Astn2* KO mice in the open field. Specifically, *Astn2* cKO mice were hyperactive and exhibited some repetitive behaviors (Supp. Fig. 4 A-C). In the three-chamber sociability test, *Astn2* cKO mice did not show differences in sociability or social novelty preference compared to the control littermates (Supp. Fig. 4 D-E). Using the rotarod, we found that *Astn2* cKO mice did not show motor coordination deficiencies. Interestingly, their performance on the rotarod was significantly better than that of control littermates (Supp. Fig. 4 F).

### *Astn2* KO animals show regionally specific changes in spine number and morphology

Different regions of the cerebellum have been implicated in coordinating specific behaviors. The hemispheres, especially Crus1, have been shown to direct social behaviors whereas posterior vermis directs inflexible and repetitive behaviors^18^. Behavior is directly correlated with synaptic connectivity in different parts of the brain. To visualize cerebellar molecular layer synapses in detail, we used Golgi-Cox staining to individually label PCs and examine their dendritic spines. We studied the numbers and subtypes of dendritic spines in three areas of the cerebellum to determine whether the loss of *Astn2* affects spines in different regions of the cerebellum differently. We were especially interested in measuring the number of immature, filopodia-like spines. We analyzed the posterior cerebellar lobule IX and crus 1. The anterior cerebellar lobule III served as a control area that has not been implicated in ASD-like behaviors but directs motor and proprioceptive functions^12^. We found no differences in spine numbers or morphology in the anterior cerebellum between wild type and *Astn2* KO littermates (Fig. 5 C). In contrast, the posterior cerebellum showed a significant increase in overall spine numbers (Fig. 5D(i)) and a significant difference in the distributions of spine lengths (Fig. 5D(ii)-(iii)). A closer look at the spine length distributions revealed a significant decrease in the fraction of long, filopodia-like spines in the *Astn2* KO animals compared to controls (Fig. 5D(iii)-(v)). Additionally, we note that though there is no statistically significant difference (p = 0.07) in the total count of filopodia (Fig. 5D(v)), we do see a sharp decrease in the average fraction of filopodia per segment (Fig. 5D(iv)). Crus 1 showed an intermediate phenotype with no differences in spine numbers (Fig. 5E(i)), but a significant difference in spine length distributions (Fig. 5E(ii)-(iii)), with a decrease in fraction (Fig. 5E(iii)-(iv)) and total count (Fig. 5E(v)) of filopodia-like spines in the *Astn2* KO animals. These data support the hypothesis that subtle changes in synaptic numbers and morphology can lead to pronounced differences in behavior.

**Figure 5:**
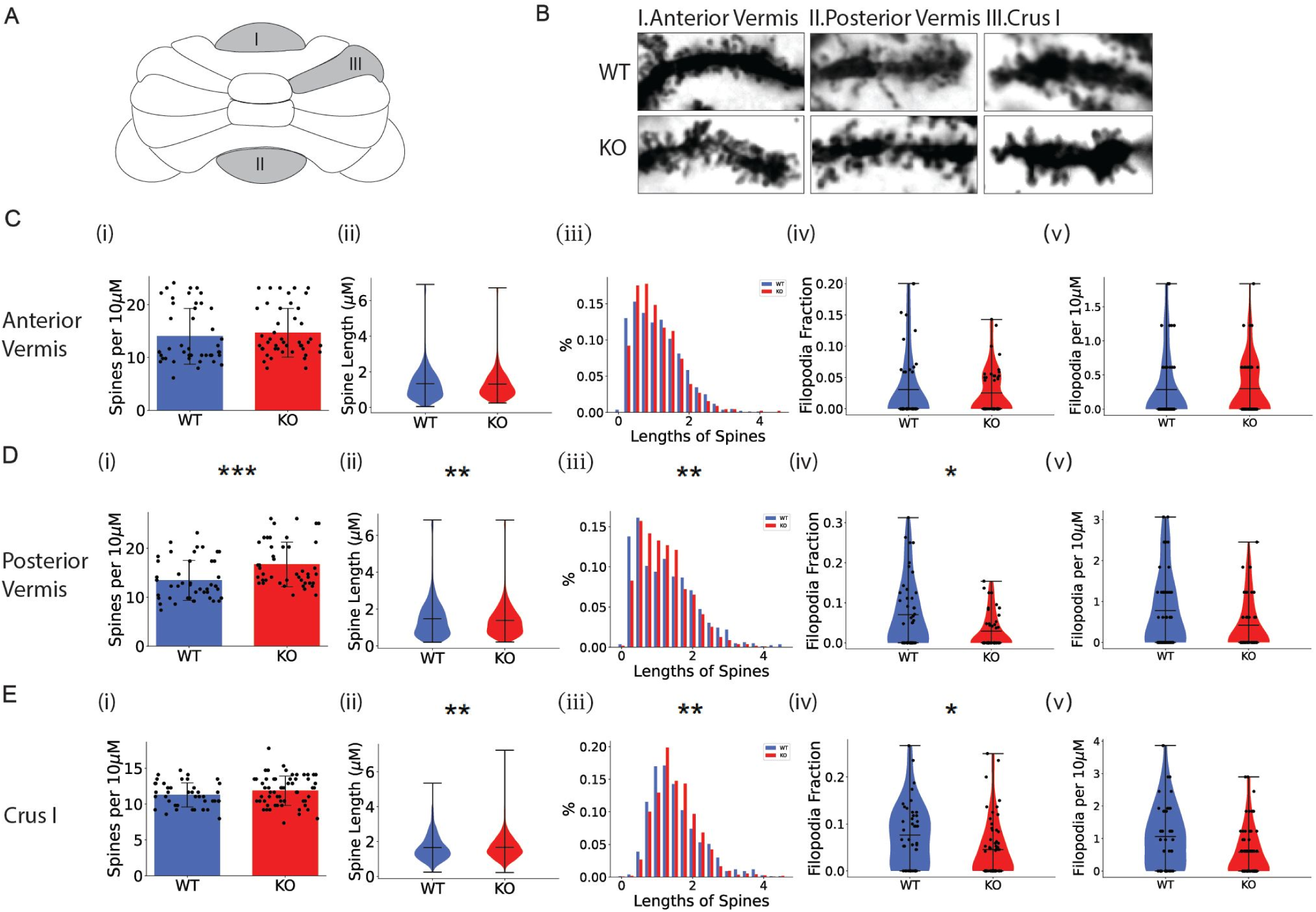
Cerebellar lobule-specific changes in Purkinje cell dendritic spine numbers and morphology in *Astn2* KO mice. **A)** Schematic of a mouse cerebellum. I marks the anterior vermis (lobule III), II marks the posterior vermis (lobule XI), and III marks Crus I. **B)** Example images of sampled dendritic segments with dendritic spines in the anterior vermis, the posterior vermis, and Crus I, in WT (*top panels*) and *Astn2* KO (*bottom panels*). **C-E)** Analysis of PC dendritic spines in WT and *Astn2* KO animals in (C) the Anterior Vermis, (D) the Posterior Vermis and (E) Crus I. **(i)** The number of spines per 10μm dendrite segment. *** p < 0.001, Wilcoxon rank-sum test. **(ii)** The distribution of spine lengths with all sampled dendritic segments pooled together. ** p < 0.01, two-sample Kolmogorov-Smirnov test. **(iii)** Detailed histogram visualization of (ii), showing the percentage proportions of each length bracket. ** p < 0.01, two-sample Kolmogorov-Smirnov test. **(iv)** Distribution of the fractions of filopodia on each sampled dendritic segment. * p < 0.05, two-sample Kolmogorov-Smirnov test. **(v)** Distribution of the total number of filopodia on each sampled dendritic segment. * p < 0.05, two-sample Kolmogorov-Smirnov test.

### *Astn2* KO animals have increased levels of the neuron-glial adhesion protein ASTN1

Since ASTN2 functions in receptor trafficking and degradation^1^, we analyzed proteomic changes that result from the loss of *Astn2*. We performed proteomic analysis on cerebellar lysates of 8 WT and 8 KO mice of both sexes. Results revealed a small number of cerebellar proteins that differed between genotypes (Fig. 6A). ASTN2 protein was highly reduced in the *Astn2* KO, as expected. *Trim32*, a small gene located within an intron of *Astn2* and transcribed from the opposite strand^8^, shows a significant increase in protein levels. RASSF8, a protein with no known function in the nervous system, was slightly reduced. Lastly, the protein levels of ASTN1, a family member of ASTN2, are highly increased in the *Astn2* KO. These results are corroborated by Western blotting (Fig. 6B). In parallel with proteomic assays, we conducted translating ribosome affinity purification (TRAP)^36,37^ followed by RNA sequencing to examine changes in gene expression after the loss of ASTN2. We crossed *Astn2* KO mice with *Pcp2-Egfp-L10a* mice expressing *EGFP*-tagged ribosomal subunit L10a under the *Pcp2* promoter to specifically profile the PC transcriptome and performed TRAP at P22. As expected, Pcp2 was highly enriched in TRAP samples, whereas the GC marker Neurod1 was enriched in input samples and depleted in TRAP samples (Suppl. Fig. 5A). In line with proteomics, differential analysis revealed only four differentially expressed genes in PCs, including *Astn2* and *Trim32* (Supp. Fig. 5 B). Input samples (whole cerebellum) revealed around 200 differentially expressed genes at a low level (Fig. 6 C), which did not result in a corresponding change in protein levels. Notably, the expression of *Astn1* mRNA was not changed in *Astn2* KO whole cerebellum or PCs (Fig. 6C, Supp. Fig. 5 C-D). These results suggest that increased ASTN1 protein levels likely result from defects in trafficking and degradation caused by the loss of ASTN2 consistent with our previous findings on the trafficking of ASTN1 by ASTN2^1,4^.

**Figure 6:**
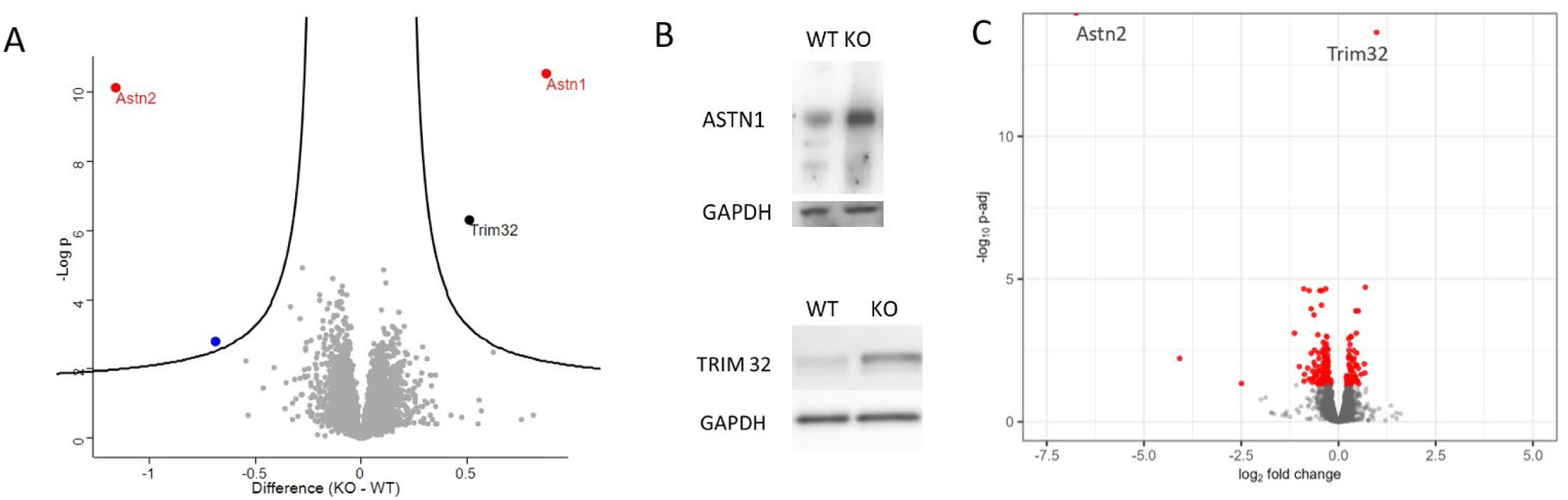
*Astn2* KO mice have increased levels of ASTN1 that is post transcriptionally regulated. **A)** Proteomic analysis of *Astn2* KO animals at P22. A volcano plot depicting differentially expressed proteins in whole cerebellar lysates of *Astn2* KO animals and wild type littermates (n = 8 WT, 8 KO). ASTN2 is downregulated and ASTN1 and TRIM32 are upregulated in *Astn2* KO animals. **B)** Western blot showing the upregulation of ASTN1 and TRIM32 in P22 cerebellar samples. **C)** Volcano plot depicting differentially expressed genes (P-adj<0.05, indicated with red) in P22 *Astn2* KO cerebellum, compared with WT littermates, identified using DESeq2. *Astn1* gene is not upregulated suggesting that ASTN1 protein overexpression is post transcriptionally regulated in *Astn2* KO animals.

### *Astn2* KO animals have an increase in Bergmann glia

Previous studies demonstrated that ASTN1 is a neuronal adhesion protein that functions in neuron-glial binding^38^ and localizes to adhesion junctions between granule cells and Bergmann glia (BG) during granule cell radial migration in development^2^. As BG form contacts with PC spines and are important for PC development and function^39,40^, we tested whether the abundance of ASTN1 protein in the *Astn2* KO might lead to changes in BG morphology or function. We used immunohistochemistry and Western blotting to examine the levels of BG marker GFAP in *Astn2* KO and WT littermates at P22. In the *Astn2* KO we observed a significant increase in the antibody signal for GFAP as well as apparent disorganization of BG fibers (Fig. 7A-B) and an increase in protein levels via Western blotting (Fig. 7C). Next, we used electron microscopy to perform an ultrastructural analysis of the interactions between BG and PCs in the molecular layer of the cerebellum. By EM, we observed an increase in BG fibers surrounding PCs (Fig. 7) in *Astn2* KO cerebellum compared to the WT. We did not observe any significant differences in the parallel fiber-PC synaptic ultrastructure between the genotypes in terms of the length of the synapse, the number of vesicles in the synaptic bouton or the contact ratio between the PF terminals and the dendritic spines (Supp. Fig. 6).

**Figure 7:**
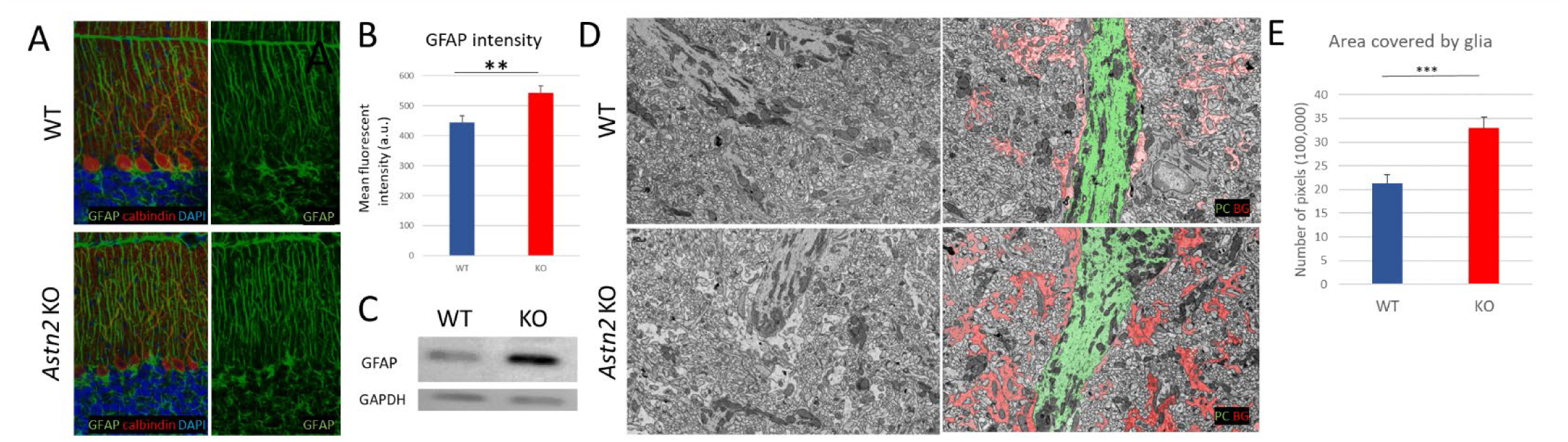
*Astn2* KO mice have an increase in Bergmann glia. **A)** Immunohistochemistry with antibodies against GFAP (a Bergmann glia marker), calbindin (a PC marker) and Hoechst in P22 *Astn2* KO mice (*bottom*) and WT littermates (*top*) (*left panels*). An increase in GFAP signal as well as disorganization of BG fibers is observed in *Astn2* KO mice (*right panels*). **B)** *Astn2* KO mice have a higher mean fluorescent intensity of GFAP staining indicating an increase in Bergmann glia. **C)** Western blot for GFAP in P22 cerebellar tissue of *Astn2* KO mice. There is an increase in amount of GFAP protein in *Astn2* KO mice. **D)** Electron microscopy imaging of WT (*top panels*) *Astn2* KO (*bottom panels*) cerebellar molecular layer at P22. An example image of 2900x direct magnification EM image (*left panel*) and a pseudocolored EM image revealing a PC dendrite (green) and Bergmann glia fibers (red) (*right panel*). **E)** Quantification of the area covered by glia in EM images in WT and *Astn2* KO animals. There is a significant increase in Bergmann glia fibers in the *Astn2* KO. 3 mice per genotype for all datasets. Data is presented as the mean + SEM. Data analyzed with Student’s T test *p < 0.05, **p < 0.01, ***p <0.001

### *Astn2* KOs show differences in Purkinje cells spontaneous and evoked synaptic currents

Given the changes in spine density and filopodial spines, we next investigated the functional properties of PC synapses in *Astn2* KO animals. Thus, we performed whole-cell electrophysiological recordings in acute brain slices from *Astn2* KO animals (P18-P25) and age-matched WT littermates. Spontaneous excitatory/inhibitory postsynaptic currents (sEPSCs/sIPSCs) were recorded to assess the overall level of synaptic input from excitatory parallel fibers (sEPSCs, Fig. 8A, *top*) and inhibitory basket and stellate cells (sIPSCs, Fig. 8B, *top*). We observed a decrease in the frequency of sEPSCs (inter-event interval, IEI, Fig. 8A, *bottom right*) as compared to WTs. We also observed significantly larger sEPSCs in *Astn2* KO animals (Fig. 8A, *bottom left*). Inhibitory currents were also altered in *Astn2* KO animals, with significantly larger sIPSCs (Figure 8B, *bottom left*) as compared to WTs. However, we did not observe a difference in sIPSC frequency (Figure 8B, *bottom right*).

**Figure 8:**
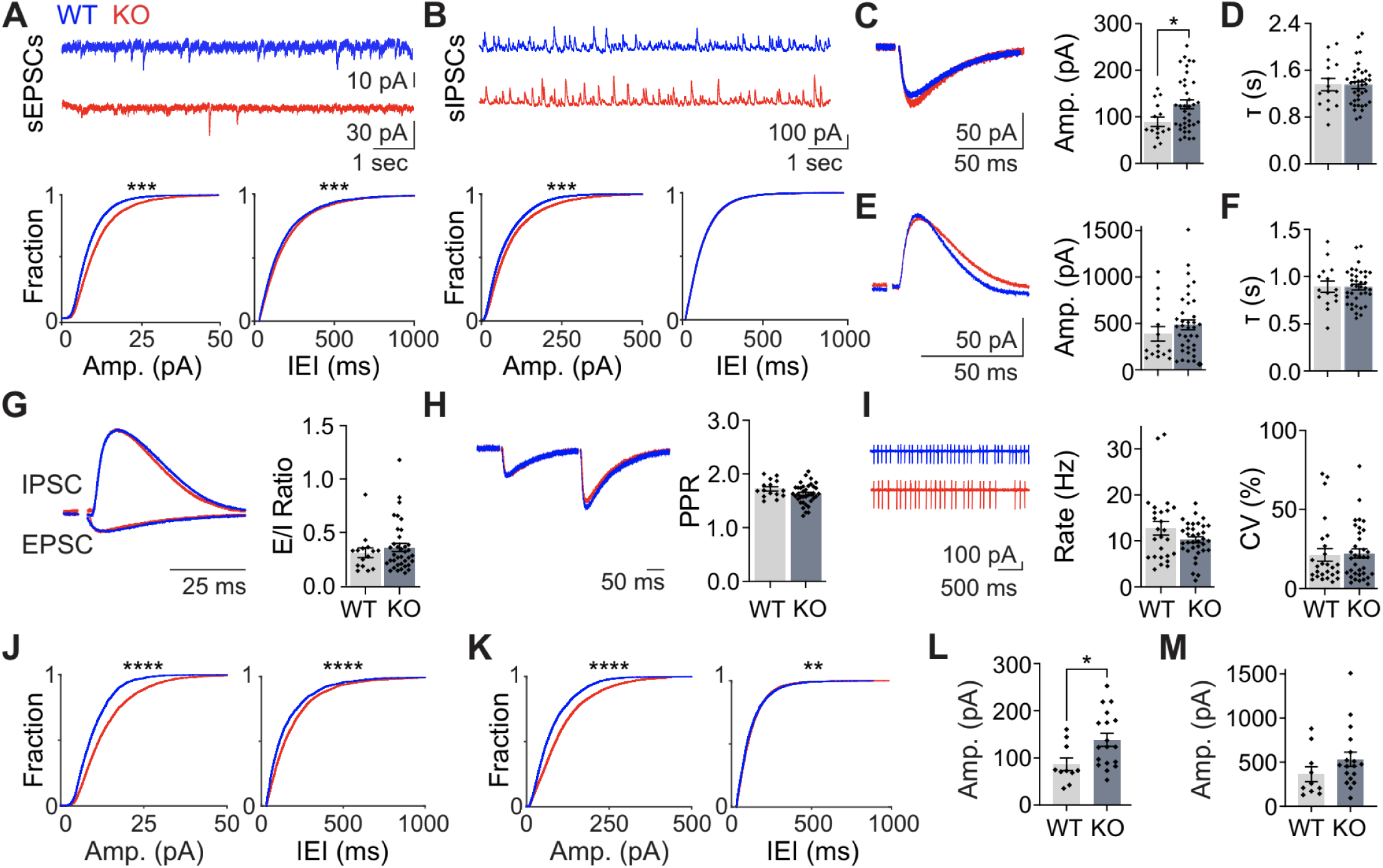
*Astn2* KO mice show differences in spontaneous and evoked synaptic currents. **A)** *Top,* spontaneous excitatory postsynaptic currents (sEPSCs) in whole-cell recordings (V_m_ = -70 mV) of Purkinje cells (PCs). *Bottom left,* cumulative distribution of sEPSC amplitudes. \*\*\**p* < 0.001, Wilcoxon rank-sum. *Bottom right,* same as *left,* for inter-event interval (IEI). \*\*\**p* < 0.001, Wilcoxon rank-sum. **B)** *Top,* spontaneous inhibitory postsynaptic currents (sIPSCs) in whole-cell recordings of PCs (V_m_ = 0 mV). *Bottom left,* cumulative distribution of sIPSC amplitudes. \*\*\**p* < 0.001, Wilcoxon rank-sum. *Bottom right,* same as *left,* for IEI. **C)** *Left,* example of mean evoked EPSC in whole-cell recordings of PCs (V_m_ = -70 mV). *Right,* summary of EPSC amplitudes across cells. \**p* < 0.05, unpaired Student’s t-test. **D)** Summary of EPSC decay kinetics (τ) across cells. **E)** *Left,* example of mean evoked IPSC in whole-cell recordings of PCs (V_m_ = 0 mV). *Right,* summary of IPSC amplitudes across cells. **F)** Summary of IPSC τ across cells. **G)** *Left,* example of mean evoked EPSC and IPSC from an individual WT and KO PC. KO traces scaled to the WT EPSC. *Right,* summary of the ratios of excitation to inhibition (E/I ratio) across cells. **H)** *Left,* example of mean evoked EPSCs from an individual WT and KO PC. KO traces scaled to the first WT EPSC. *Right,* summary of paired-pulse ratios (PPR) across cells. **I)** *Left,* example of mean extracellular recordings of PC spiking activity from an individual WT and KO PC. *Middle,* summary of spike rates across cells. *Right,* summary of coefficients of variation (CV) across cells. **J)** *Left,* cumulative distribution of sEPSC amplitudes of PCs in posterior vermis. \*\*\*\**p* < 0.0001, Wilcoxon rank-sum. *Right,* same as *left,* for IEIs. \*\*\*\**p* < 0.0001, Wilcoxon rank-sum. **K)** *Left,* cumulative distribution of sIPSC amplitudes of PCs in posterior vermis. ****p < 0.0001, Wilcoxon rank-sum. *Right,* same as *left,* for IEIs. \*\**p* < 0.01, Wilcoxon rank-sum. **L)** Summary of EPSC amplitudes across PCs in posterior vermis. \**p* < 0.05, unpaired Student’s t-test. **M)** Summary of IPSC amplitudes across PCs in posterior vermis. Error bars ±SEM.

We next recorded evoked EPSCs from parallel fibers (Fig. 8C, *left*) and found that *Astn2* KOs have greater EPSC amplitudes as compared to WTs (Fig. 8C, *right*). Similarly, we observed a trend toward larger evoked IPSCs (Fig. 8E). These results are consistent with our observations of larger sEPSCs and sIPSCs and may reflect an enhanced postsynaptic response. We did not, however, observe any difference in EPSC or IPSC kinetics (Fig. 8D,F), suggesting no difference in average synapse location on the somatodendritic axis of PCs, or change in the receptor subunit compositions between WT and *Astn2* KO animals.

Changes in both spontaneous and evoked EPSCs in the *Astn2* KOs could alter the balance of excitation and inhibition, as a bias toward excitation has been linked to cognitive disorders such as ASD^41–44^. We therefore also calculated the ratio of excitation to inhibition (E/I ratio, Fig. 8 G, *left*) but found no difference in E/I ratio between *Astn2* KOs and WTs (Fig. 8G, *right*), consistent with our observation that both spontaneous EPSCs and IPSCs (Fig. 8A,B) and evoked EPSCs and IPSCs (Fig. 8C,E) increase in tandem with a global knockout of *Astn2*.

Supporting the interpretation that changes in synaptic amplitude are due to postsynaptic effects, we found that the paired-pulse ratio of EPSCs onto PCs (PPR, Fig. 8H, *left*) was unchanged in *Astn2* KO animals (Fig. 8H, *right*), suggesting no difference in presynaptic release probability. Finally, we measured the intrinsic excitability of PCs using noninvasive cell-attached recordings (Fig. 8I, *left*) and found no difference in either spiking rate (Hz, Fig. 8I, *middle*) or coefficient of variation (CV, Fig. 8I, *right*).

Notably, our recordings were performed in both the anterior and posterior vermis of the cerebellum. Because anatomical differences were most pronounced in the posterior cerebellum, we next subdivided our recordings to specifically assess synaptic responses in this region. Importantly, changes in synaptic transmission were more pronounced in the posterior vermis, with larger differences in both sEPSC amplitude (Fig. 8J, *left*) and frequency (Fig. 8J, *right*), as well as sIPSC amplitude (Fig. 8K, *left*). In this region, we also measured a small but statistically significant reduction in sIPSC frequency (Fig. 8K, *right*). PCs in the posterior vermis showed larger evoked EPSC amplitudes (Fig. 8L) and trend towards larger evoked IPSC amplitudes (Fig. 8M). Together, these measurements suggest both pre- and postsynaptic changes in synaptic transmission. Specifically, the decrease in sEPSC and sIPSC rate suggests fewer functional excitatory and inhibitory synapses onto KO PCs. While our anatomical measurements indicate greater spine numbers, these results may be related to the reduced fraction of filipodial spines in KO animals, or single parallel fibers impinging on more spines (fewer unique contacts). In contrast, the significantly larger sEPSC and sIPSC amplitudes may suggest a postsynaptic enhancement of synaptic transmission, perhaps reflecting larger numbers of postsynaptic AMPA and GABAA receptors at mature spines in KO animals. Overall, these results show subtle but significant changes in the synaptic strength of PCs, with a stronger effect on sEPSCs, suggesting that cerebellar processing is altered by *Astn2* loss of function.

## DISCUSSION

Our studies of mice lacking *Astn2* demonstrate an important role for ASTN2 in ASD-related behaviors including communication via USVs, social behavior, hyperactivity, repetitive behaviors, and in cerebellar circuit properties. In addition, mutant animals appear to have reduced cerebellar learning as measured by eyeblink conditioning. Proteomic analyses and ultrastructural studies showing an increase in BG volume surrounding PC spines are consistent with changes in neuron-glial interactions. A change in the number and maturity of spines in the posterior vermis and Crus 1 supports the interpretation that ASTN2 is important for spine formation and dynamics in areas of the cerebellum associated with social and repetitive behaviors. Electrophysiological studies show changes in the *Astn2* KO cerebellum, especially in its posterior aspect, consistent with the results on spine changes in this area. In agreement with other studies^24,25^ our studies support the conclusion that subtle changes in cerebellar anatomy and physiology can lead to significant changes in behavior.

As ASDs are characterized by social impairments, repetitive and inflexible behaviors^21,25^, and in many cases, language deficits^1,45^, we focused on analyzing changes in these behaviors in *Astn2* KO mice. While human language cannot be explored in mice, vocal communication behavior is conserved across taxa^46^. Our assays of USV calls in a pup separation assay showed a dramatic (40%) decrease in calls and changes in call structure. These findings represent one of the largest decreases in pup isolation calls in mouse models of ASD and are consistent with prior studies on children with *ASTN2* CNVs^1^ as well as other studies in mouse models^27,28,47^. Shorter duration of calls with an unmatched increase in pause duration and a comparable peak amplitude of the sound pressure level, or volume, of the calls that we observed suggest that the reduction in KOs is not due to respiratory difficulties in holding the call. Spectrally, the comparable average pitch frequency of the calls indicates that abnormalities in the laryngeal musculature is likely not driving the call reduction, as the KO mice could reach a similar pitch. The narrowed pitch range and reduced fraction of dynamic calls suggests a less comprehensive composition of syllables among the KO calls compared to those from control littermates. Thus, our analysis reveals early social communicative challenges with the disruption of *Astn2*. As the *Pcp2-Cre* line we used to generate a cKO specific to cerebellar PCs is not expressed until the end of the first postnatal week, we were unable to assay USV call frequency and structure in pups with a PC-specific loss of *Astn2*.

Our findings show that *Astn2* KO animals have impaired social behavior, as measured in the three-chamber tests for sociability and social novelty preference. *Astn2* KO mice exhibited relatively normal sociability as they showed a preference for the social versus a non-social stimulus. Nevertheless, they spent overall less absolute time with the social stimulus compared to the control animals. Importantly, *Astn2* KO mice showed no preference for social novelty as they spent the same amount of time with a familiar mouse as with the novel mouse in contrast to controls. This suggests that ASTN2 is important for influencing social behavior. Interestingly, PC-specific *Astn2* cKO animals did not show impairments in social behavior in a three-chamber paradigm, suggesting that other cell types or brain regions contribute. Recently, a study has reported differences in the spine number, layer thickness and numbers of neuronal cell bodies in the hippocampus and prefrontal cortex in *Astn2* KO animals, which may partially explain the differences observed in social behavior^48^. More research is needed to elucidate which brain regions contribute to deficits in social behavior in *Astn2* KO animals and the exact contribution of the cerebellum to social behavior, language, and communication.

*Astn2* KO animals have a significant increase in hyperactivity and repetitive behaviors as seen in the open field assay. Repetitive behaviors are a hallmark of ASD and hyperactivity is a very common comorbidity. In addition, we found that global *Astn2* KOs have a reduced anxiety phenotype in the open field. A similar result has been recently reported in *Astn2* KOs using an elevated plus-maze test^48^. Importantly, we found that PC-specific cKO mice have a similar phenotype to global KO animals in hyperactivity and repetitive behaviors, suggesting that the cerebellum plays an important role in regulating those behaviors.

The fact that we did not observe changes in social behavior in *Astn2* cKO mice is consistent with our anatomical studies showing no, and less pronounced, differences in Crus 1 of the cerebellum in spine numbers and spine morphology, respectively. In comparison, the posterior vermis, which is associated with repetitive behaviors^18^, has significant differences in spine and filopodia numbers that correlate well with repetitive behavioral changes seen in *Astn2* cKO. Interestingly, we did not observe any differences in *Astn2* KO motor reflexes and coordination. *Astn2* KO mice could right themselves as quickly as controls as pups and performed equally well on the rotarod as adults. Many animal models that focus on the cerebellum and ASD show motor differences, but some do not (for a comprehensive list see ^49^). We hypothesize that the localized differences in spine numbers and morphology and the lack of such differences in the anterior lobule of the cerebellum where many motor behaviors are regulated might explain this phenotype. As anatomical and developmental studies did not reveal any significant changes in neurogenesis, glial-guided migration or formation of the neuronal layers (Supp. Fig. 2,3), it is unlikely that the behavioral changes were due to defects in basic steps in cerebellar development but rather subtle differences at the level of the synapse.

The molecular analyses we carried out on the *Astn2* KO cerebellum indicated that the primary change after the loss of *Astn2* was a dramatic increase in the protein levels of the neuron-glial ligand ASTN1. This finding is consistent with our prior studies showing that ASTN2 binds to and promotes the trafficking of ASTN1^1,4^. Prior studies on ASTN1 demonstrated that it functions in neuron-glial adhesion^38^ by binding N-cadherin on BG fibers during granule cell radial migration in development^3,50,51^. In support of changes in neuron-glial binding in the mutant cerebellum, Western blotting and immunohistochemistry showed an increase in GFAP protein levels and EM analysis revealed a large increase in the volume of BG processes around PCs in the molecular layer. BG, PCs and granule cells form a glutamatergic tripartite synapse in the cerebellar cortex^52^. There is a very intimate relationship between PCs and BG both during development, where they influence each other’s growth and maturation^39,40^, as well as in adulthood, where BG has a role in synaptic pruning^53^ and plasticity^54^. BG ensheath PC dendritic spines, which is important for proper functioning of the synapses^55^. This raises the possibility that the loss of *Astn2* leads to changes in the stability or plasticity of dendritic spines in mutant animals. Golgi analyses of dendritic spines indeed demonstrated changes in the number and maturation state of spines in mutant animals as they showed an increase in spine number and a reduction in the fraction of filopodial protrusions, considered an immature state of spine maturation^56^. It was of special interest that the spine changes we observed were primarily in the posterior vermis, an area associated with inflexible/repetitive behaviors^18^. We hypothesize that the increase in BG area as a direct result of the increase in ASTN1 protein levels leads to the increased spine density and disrupted synapse maturation in *Astn2* KO mice. More research is needed to validate this hypothesis.

In line with the observed changes in spine density and maturation, we also found corresponding differences in synaptic transmission in *Astn2* KO animals. Specifically, we observed reduced frequencies of spontaneous excitation and inhibition, and increased amplitudes of both spontaneous and evoked excitation and inhibition. Notably, these effects were preferential to the posterior vermis, where anatomical differences were observed. It is somewhat surprising that we observed decreased rates of spontaneous synaptic currents when anatomical measurements showed an overall increase in spine number. However, one possible explanation could be that the increased number of mature spines is also associated with a redundant sampling of parallel fiber inputs (i.e. fewer unique inputs). In agreement with our measurements, such a configuration would result in both the reduced frequency and increased amplitude of events, as the redundant inputs would boost the amplitude of action potential-evoked EPSCs and IPSCs. Fully resolving this question would require reconstruction of individual parallel fibers and represent an interesting future direction.

Our loss of function *Astn2* studies are consistent with our prior work showing that ASTN2 regulates synaptic protein receptor trafficking and PC synaptic properties^1^. As receptor trafficking is critical for synaptic plasticity and function, these findings are also consistent with prior studies implicating defects in pre- and postsynaptic synaptic proteins (e.g. Neurexins, Neuroligins, Synapsin 1 and 2, PSD-95, Cadherins, and Protocadherins, Shank3) ^29,57–59^ in ASD-like behaviors. Our studies on ASTN2, therefore, support the model that changes in circuit properties contribute to ASD-like behaviors. Importantly, the changes in synaptic transmission we observed were relatively subtle. This may explain the lack of obvious motor deficits in these animals and support the idea that even small changes in cerebellar processing can affect cognitive behaviors. Moreover, the preferential deficits we observed in the posterior cerebellum are in line with the suggested role of this area in ASD-like behaviors, and particularly repetitive behaviors. Therefore, our analyses of *Astn2* KO and cKO mice reveal an important role for ASTN2 in ASD-related behaviors and cerebellar circuit properties.

## ACKNOWLEDGEMENTS

We are grateful to Dr. Erich Jarvis for providing access to USV testing equipment and to Dr. Nathaniel Heintz for providing access to a three-chamber assay system, open field testing device and the rotarod. We thank Dr. Carol A. Mason for comments on the EM experiments. We thank Joe Rodriguez and Eve Govek for help with sample preparation for EM experiments. The Gensat Project provided the *Pcp2-Cre* line used to generate the cKO animals. We thank Drs. Matt Paul and Tom Carroll from the Bioinformatics Resource Center at Rockefeller University for their support with bioinformatics analysis. We thank the Bioimaging Centre at Rockefeller University for its support with imaging and image analysis. Figures created using BioRender.com.

Supported by NIH grant 5R01NS116089 (MEH and CH); NIH grants R01NS128054 and R01NS112917 (CH); Fellowships from the Sigrid Juselius Foundation and the Leon Levy Foundation (KM); grant # UL1 TR000043 from the National Center for Advancing Translational Sciences (NCATS, National Institutes of Health (NIH) Clinical and Translational Science Award (CTSA) program (MH) and and NICHD grant P50HD103525 to IDDRC@WUSTL (SEM).

## AUTHOR CONTRIBUTIONS

MH carried out all developmental studies, anatomical studies, behavioral tests including USV recordings, three-chamber, open field and rotarod tests, Golgi studies of spine structure, EM studies, proteomic studies and writing the manuscript. KF carried out electrophysiology and eyeblink conditioning experiments and analysis. SEM and JDD carried out the analyses of USV datasets, and SEM manuscript writing. SG helped make the *Astn2* KO and cKO floxed constructs and the KO mutant and cKO animals. HAP carried out the EM. KM carried out TRAP and RNA sequencing. SH carried out the proteomic experiments and analysis. JZ carried out the imaging and analysis of Golgi studies of PC spines. CH planned and carried out the electrophysiology and eyeblink experiments and writing the manuscript. MEH assisted with planning the behavior, anatomical, spine studies, EM and molecular studies and writing the manuscript. All authors read and approved the manuscript.

## METHODS

### Mice

All procedures were performed according to the guidelines approved by The Rockefeller University Institutional Animal Care and Use Committee. Males and females were used for all studies and were randomly allocated to groups. Pcp2-cre line mice were obtained from the GENSAT project. Tg(Pcp2-Egfp-L10a) TRAP mice (RRID: IMSR_JAX:030267)^36^ were a gift from Dr. Nathaniel Heintz. For electrophysiology and eyeblink conditioning experiments, all animal care and experimental procedures were approved by the Duke University Institutional Animal Care and Use Committee. Experiments were conducted during the light cycle with both male and female mice. adult. All mice were housed with 12-hour light/dark cycles with food and water ad libitum.

### Generation of Astn2 knockout lines

C57BL/6J mice (The Jackson Laboratory) were used to create the *Astn2* KO and *Astn2^flox/flox^* mice. To knock out Astn2, two guide RNAs flanking the promoter and the exon containing ATG start codon were selected. RNP containing guide RNA and cas9 protein was co-injected into C57BL/6 fertilized embryos at the Rockefeller University transgenic core facility. To assess the deletion of Astn2, genomic DNA from founders was isolated using the alkaline lysis method. Primers flanking the outside of the CRISPR sgRNAs target sequences were designed and amplified with Taq polymerase. The PCR products from the positive founders were sequenced to confirm the deletion and the locus integrity. Non-specific integration was characterized by two simple PCR amplifications. The loss of ASTN2 protein expression was confirmed using Western blot analysis and proteomic analysis of whole cerebellar tissue. Astn2 KO mice did not show reduced survival or fertility and their weights were comparable across genotypes at P22.

To generate the *Astn2^flox/flox^* mouse line, two sgRNAs targeting the exon containing ATG start codon were designed and made. A donor plasmid containing the exon flanked by two loxP sites and two homology arms of 800 bp each was designed and constructed. Briefly, RNP containing two sgRNAs and Cas9 protein was prepared. Pronuclear injections of RNP with the 10ng/μl of donor DNA was performed at the Rockefeller University transgenic core facility with embryos from C57BL/6. Genotyping with primers flanking the two loxP sites for potential founder mice were performed as described above. PCR products were analyzed by Sanger sequencing to validate the 5′ and 3′ loxP insertions in the mouse genome. The *Pcp2-Cre* line (Gensat) was then crossed with the floxed line to generate a Purkinje cell specific conditional loss of function line.

### Immunohistochemistry

Briefly, 100um vibratome sections were blocked with 15% normal horse serum (Gibco), 0.3% Triton in PBS overnight and then incubated with primary antibodies overnight at 4°C, washed in PBS 3x 15 mins and incubated with Alexa Fluor® secondary antibodies overnight at 4°C. Sections were mounted with ProLong® Gold anti-fade mounting media and covered with 1.5 thickness Fisherbrand cover glass.

### Antibodies used for immunohistochemistry (IHC) and Western blot (WB)

Primary antibodies, anti-ASTN2 (1:500 for WB, custom made), anti-Calbindin D28-k (1:500 for IHC, Swant #CB38), anti-GFAP (1:2000 for IHC, 1:1000 for WB, Dako #Z0334), anti-ASTN1 (1:1000 for WB, custom made), anti-TRIM32 (1:500, ThermoFisher #10326-1-AP) anti-GAPDH (1:10,000 for WB, Chemicon #mab374), anti-PH3 (1:100 for IHC, Cell Signalling #53348), Hoersht (1:10000 for IHC, ThermoFisher #62249). Secondary antibodies: Donkey anti-mouse, -rabbit, and -goat IgG conjugated to Alexa 405, 555, 633, and 647 (abcam) all used at 1:500. HRP-conjugated secondary antibodies (Jackson Immunoresearch) were used at 1:8000 (anti-mouse, # 515-035-062) or 1:3000 (anti-rabbit #111-035-144) for Western blots.

### Imaging

Immunohistochemistry images were acquired using an inverted Zeiss LSM 880 NLO laser scanning confocal microscope with a Plan-Apochromat 20x/1.4 NA objective lens and 2x digital zoom. Images were acquired by setting the same gain and offset thresholds for all images per experiment and over/underexposure of signal was avoided. Images were quantified in FIJI/ImageJ (version 1.53c).

### Maternal isolation-induced ultrasonic vocalizations

During the first two weeks postnatal, we assessed the *Astn2* WT, Het and KO littermates for signs of gross developmental and communicative delay following our previously published methods ^65,66^. Isolation from the dam induces ultrasonic vocalizations (USVs) in the mouse pup to elicit maternal care, which is one of the earliest forms of social communication we can examine in the mouse. This behavior also has a developmental trajectory as it begins just after birth, peaks during the first week postnatal and then disappears around P14, making it useful for examining delay in early social circuits. USVs were recorded at P5 - P14 following our previously published methods ^65,67^. Briefly, the pup was removed from the nest and placed in an empty beach cooler (internal dimensions are L 27 x W 23 x H 47 cm) that acts as a sound attenuation box studio. USVs were recorded for five minutes using an Avisoft UltraSoundGate CM16 microphone, Avisoft UltraSoundGate 116H amplifier, and Avisoft Recorder software (gain = 5 dB, 16 bits, sampling rate = 250 kHz). The pup was then weighed and returned to the nest. Frequency sonograms were prepared from raw WAV recordings in MATLAB (frequency range = 25 kHz to 120 kHz, FFT size = 512, overlap = 50%, time resolution = 1.024 ms, frequency resolution = 488.2 Hz). Individual syllables and other temporal and spectral features were identified and counted from the sonograms using a custom MATLAB pipeline as previously described ^68,69^. USV analysis was conducted using SPSS Statistics v29. Linear mixed models were used to analyze the USV data with genotype as a fixed predictor and age as a random, repeated factor, nested in litter and subject. Simple main effects were used to dissect significant genotype*age interactions. Sex was a non-significant predictor and was removed to achieve the most parsimonious models. Several variables, including number of calls, bouts of calls, pitch range of flat and dynamic calls, average pitch of dynamic calls, and average pause within call bouts, were square root transformed due to non-normal distributions to meet the assumptions of parametric testing.

### Behavioral experiments

The testing apparatuses were cleaned with 70% ethanol and water between test runs and with Clydox when switching between genders. The experimenter was blind to the treatment for each group. In all experiments, mice were habituated to the testing room for at least 1-hour prior to testing and were run during their light cycle. All experimental groups were run during the same session to exclude batch effects. Assays were performed by examiner blinded to genotype.

### Three chamber social behavior apparatus

Animals were tested between 8-12 weeks of age. Animals were tested in the three chambered apparatus as previously described^70^. Briefly, mice were allowed to freely explore 3 connected 50 x 50 x 30 cm open-top plexiglass containers. A cohort of wild type males and females, age matched to the experimental group, were habituated to a wire cup for 15 min daily, three days prior to testing. On testing day, mice from the experimental groups were habituated to the arena with wire cups for 10 mins. In phase two of the task, a novel, conspecific wild type mouse of the same sex was placed under a wire cup in one of the outer chambers, while a plastic lego block was placed under a wire cup in another outer chamber. Gates were removed allowing free access of the experimental mouse to both adjacent chambers for 10 minutes. In phase three of the task, the lego was replaced with a novel, conspecific wildtype mouse of the same sex, whereas the previously run conspecific mouse is classified as the familiar mouse. The experimental mouse was reintroduced to the center chamber, and gates were removed allowing free access to investigate the novel and familiar mouse for 10 minutes. Time spent interacting with the animals and objects (Lego block) was recorded by the examiner with a stopwatch.

### Open field locomotion

In the open field test, 8-12 week old mice were lowered into and allowed to explore a clear plexiglass 50 x 50 x 30 cm open-top cube for 1 hour. The arena was illuminated to 230 lux. Up to 8 mice per trial were run in separate arenas. Mouse location, vertical rears, and velocity was automatically tracked by Fusion Software. A zone map delineating the center zone (4×4) and the periphery zone (7-4 x 7-4) was used to determine duration in the center and periphery.

### Light/dark box

The light/dark box consisted of a black acrylic apparatus, partitioned into an exposed (light, 230 lux) and closed (dark, 0 lux) compartment, placed in a clear plexiglass 50 x 50 x 30 cm open-top cube. Mice were lowered into the dark partition and their location and velocity was tracked automatically using Fusion Software. A zone map partitioning the exposed and close compartment was used to track duration in both compartments and latency to enter the exposed compartment.

### Righting reflex

P7 pups were removed from their cage during their awake time and placed in a supine position. The time to completely right themselves was measured with a stopwatch by the experimenter. Pups were returned to the cage immediately.

### Rotarod

The rotarod machine used is the ENV-575M by Med Associates Inc. The accelerating speed mode was used where the drum begins at the lowest speed (4RPM) and takes five minutes to steadily reach the highest speed (40RPM). Five animals of random genotype were tested at the same time in five zones. Each zone has a photo beam that senses when the animal falls off the treadmill and automatically records the breaking of the beam. Animals were tested on five consecutive days. The average latency to fall of three trails per day was calculated.

### Surgical procedures for delay eyeblink conditioning

All surgeries were performed under anesthesia using an initial dose of ketamine/xylazine (50 mg/kg) five minutes before and 1-2% isoflurane throughout surgery. Mice received an initial dose of meloxicam (5 mg/kg). Breathing rate and toe pinch responsivity were continuously monitored during surgeries. A heating pad (TC-111 CWE) was used to maintain body temperature. Titanium headplates (HE Parmer) were attached to the skull with Metabond (Parkell). Mice received buprenex (0.05 mg/kg) and cefazolin (50 mg/kg) every 12 hours for 48 hours after surgery. Mice were given one week for recovery before habituation to head fixation and training.

### Delay eyeblink conditioning

The behavioral setup was constructed according to Heiney et al., 2014^35^. Prior to experiments, all mice were habituated to head fixation on the same cylindrical treadmill used for training for 60 min/day until they could walk comfortably on the wheel with no signs of distress (3-5 days). Stimulus delivery and frame acquisition for video monitoring were triggered with an Arduino Uno microcontroller board (Arduino) controlled by modified Arduino and MATLAB (MathWorks) code written for Neuroblinks software^35^. Mice were trained during daily sessions of 200 trials in which a 30-ms air puff (30 PSI) was delivered 3 mm from the mouse’s right lateral cornea (unconditioned stimulus, US) and paired with a co-terminating, 250-ms blue LED (conditioned stimulus, CS) positioned 10 cm in front of the mouse with a minimum inter-trial interval of 15 seconds. Five percent of all trials contained CS-only probe trials where the US was omitted, and three percent of all trials contained US-only test trials. Trials were only initiated if the eyelid was at least 80% open for at least 200 ms. A CR was counted if the eyelid was at least 10% closed in the last 30 ms before US delivery. Criteria were occasionally relaxed during periods of squinting.

### Proteomics

P21 cerebella were lysed using lysis buffer (50mM PBS, pH 7.4, 150mM sodium chloride, 0.5 % sodium deoxycholate, 1% NP-40, and 1X protease inhibitor cocktail). Proteins were dissolved in 8M urea, 50mM triethylammonium bicarbonate (TEAB). Reduction and alkylation of cysteines was carried out using dithiothreitol (DTT) and iodoacetamide (IAA). Proteins were extracted using chloroform/water/methanol extraction and pellets were dissolved in 100mM TEAB containing endoproteinase Lys-C. Digestion was carried out overnight at room temperature followed by the addition sequencing-grade trypsin and further digesting for 6 hours at 37°C. Peptides were labeled with TMTpro mass tag according to manufacturer specifications. The labeling efficiency was evaluated by pooling aliquots from all channels, storing the remaining material at -80°C meanwhile. Upon verification of labeling efficiency and stoichiometry, reactions were quenched using hydroxylamine and samples were pooled. Pooled peptides were fractionated using a high-pH spin column into 8 fractions. The fractions were analyzed by LC-MSMS using an Easy nLC1200 equipped with a 250mm*75µm Easyspray C18 column. Peptides were separated across a 120-minute gradient and analyzed by an orbitrap Fusion Lumos mass spectrometer operating in positive-ion mode. Fragmentation was done in an MS2-data-dependent fashion. Spectra were queried against the Mus musculus proteome at 1% FDR. Further quantitation was performed within the Perseus computational environment.

### Translating ribosome affinity purification (TRAP)

TRAP was performed as previously described^36,37^. Briefly, Streptavidin MyOne T1 Dynabeads (Invitrogen 65601) were coated with biotinylated protein L (Pierce 29997) for 35 min at room temperature, washed with 3% IgG and Protease-free BSA (JacksonImmuno 001-000-162) in 1× PBS and subsequently incubated with 50 µg each of 19C8 and 19F7 anti-GFP monoclonal antibodies (Memorial Sloan-Kettering Monoclonal Antibody Facility) in 0.15 M KCl TRAP wash buffer (10 mM HEPES-KOH at pH 7.4, 5 mM MgCl2, 150 mM KCl, 1% NP-40, supplemented with 100 µg/mL cycloheximide [Millipore Sigma C7698-1g] in methanol, 0.5 mM DTT [Thermo Fisher Scientific R0861], 20 U/mL RNasin [Fisher Scientific PR-N2515]) for 30 min at room temperature using end-over-end rotation. After antibody binding, the beads were washed three times with 0.15 M KCl TRAP wash buffer and resuspended in 0.15 M KCl TRAP wash buffer. Each reaction was supplemented with 30 mM DHPC (Avanti 850306P). Cerebella from P21 Astn2 wild-type and knockout mice crossed with Tg(Pcp2-Egfp-L10a) mice were dissected and placed in TRAP dissection buffer (2.5 mM HEPES-KOH at pH 7.4, 35 mM glucose, 4 mM NaHCO3 in 1× HBSS, supplemented with 100 µg/mL cycloheximide). The tissue was homogenized in chilled TRAP lysis buffer (10 mM HEPES-KOH at pH 7.4, 5 mM MgCl2, 150 mM KCl, supplemented with 0.5 mM DTT, 100 µg/mL cycloheximide, protease inhibitor cocktail [Sigma 11836170001], 40 U/mL RNasin, and 20 U/mL Superasin [Thermo Fisher Scientific AM2694]). The homogenate was centrifuged at 2000g for 10 min at 4°C. 3% of the supernatant was set aside as an input. The rest of the supernatant was supplemented with NP-40 to a final concentration of 1% and with DHPC to a final concentration of 30 mM and incubated for 5 min on ice. The samples were centrifuged at 20,000g for 10 min at 4°C, and the supernatant was used for immunoprecipitation with GFP-conjugated beads overnight at 4°C with end-over-end rotation. Subsequently, the beads were washed four times in 0.35 mM KCl TRAP wash buffer (10 mM HEPES-KOH at pH 7.4, 5 mM MgCl2, 350 mM KCl, 1% NP-40, supplemented with 100 µg/mL cycloheximide, 0.5 mM DTT, and 20 U/mL RNasin) and resuspended in 100 µL of RLT buffer from the RNeasy Micro Kit (Qiagen 74004) supplemented with 1% β-mercaptoethanol. The resuspended beads were incubated at room temperature for 10 min, placed on a magnet, and the supernatant containing the RNA was collected and purified using the RNeasy Micro Kit, including dsDNAse treatment. TRAP efficiency was determined using Pcp2 and Neurod1 qPCR (not shown). RNA integrity was determined using Bioanalyzer RNA 6000 Pico kit (Agilent) prior to RNAseq library preparation.

### RNA sequencing

RNA-seq libraries were prepared from 100 ng RNA/sample using the NEBNext Ultra II RNA library preparation kit for Illumina (NEB E7770S) in conjunction with the NEBNext poly(A) mRNA magnetic isolation module (NEB E7490) and NEBNext multiplex oligos for Illumina (NEB E7335 and E7500). The quality of the sequencing libraries was evaluated using the Agilent 2200 TapeStation with D1000 High-Sensitivity ScreenTape. The samples were sequenced at the Rockefeller University Genomics Resource Center using NovaSeq 6000 (Illumina) to obtain 75-bp single-end reads. Samples with a sufficient number of reads after QC were included in the analysis.

### Bioinformatics analysis for RNA-seq

Sequence and transcript coordinates for mouse mm10 UCSC genome and gene models were retrieved from the Bioconductor Bsgenome.Mmusculus.UCSC.mm10 (version 1.4.0) and TxDb.Mmusculus.UCSC.mm10. knownGene (version 3.4.0) libraries, respectively. RNAseq reads were mapped using Rsubread (version 1.30.6)^60^. Transcript expressions were calculated using Salmon (version 0.8.2)^61^, and gene expression levels as TPMs and counts were retrieved using Tximport (version 1.8.0)^62^. Normalization of raw read counts in genes and differential gene expression analysis were performed using DESeq2 (version 1.34.0)^62^. Gene expression was considered significantly different when P-adj < 0.05. Normalized, fragment-extended signal bigWigs were created using rtracklayer (version 1.40.6)^63^ and visualized in Integrative Genomics Viewer (IGV)^64^.

### Golgi staining and PC dendritic spine analysis

P22 cerebella were processed for Golgi staining using the FD Rapid GolgiStain™ Kit. During processing 100um sections were cut using Laica vibratome (VT1200S; Leica) and mounted with Permount® mounting medium. Slices were imaged using an upright wide field brightfield Axioplan 2 microscope fitted with a 40x or 63x objective (Zeiss).A total of 3-4 P21-22 mice were imaged per genotype. For each lobule in our analysis (posterior vermis, anterior vermis, and Crus I), a total of 772 to 1160 spines per genotype per lobule are included in our analysis.

These are sampled from 4-5 PCs per lobule per mouse, with 3 dendritic segments from each PC. Each dendritic segment is either 10μm (when using 63x magnification lens) or 16 μm (when using 40x magnification lens) in length. These segments are normalized when pooled together for statistical analysis. Length of a spine was measured from the edge of the dendritic shaft until the end of the spine length. Spines longer than 2.773 μm were characterized as filopodia. All analyses on PC dendritic spines are performed using custom scripts in Python.

### Electron Microscopy

Mice were perfused transcardially with 2% glutaraldehyde and 2% paraformaldehyde in 0.1 M sodium cacodylate buffer (pH 7.2). Sagittal slices (100 um thickness) were cut through the cerebellum with a vibratome. They were post-fixed for 2 hours with 1% osmium tetroxide/1.25% potassium ferrocyanide in 0.1 M sodium cacodylate buffer (pH 7.2), stained with 0.5% uranyl acetate in 0.05 M maleate buffer pH 5.2, dehydrated with graded acetone, and embedded in Eponate 12 (Ted Pella, Inc). Ultrathin sections (60–65 nm) were then counterstained with uranyl acetate and lead citrate, and images were acquired using a Tecnai Spirit transmission electron microscope (FEI, Hillsboro, Oregon) equipped with an AMT BioSprint29 digital camera.

### EM image analysis

20 images were taken in molecular layer of Crus I (2900x magnification) from 3 mice per genotype and analyzed in Adobe Photoshop. Osmium fixation used enhances the membranes and gives the Purkinje cell cytoplasm a darker appearance, while Bergmann glia are whitish pale. Purkinje processes are rich in dark mitochondria and stacks of smooth ER, while Bergmann glia is not. We used the Magic Wand function in A dobe Photoshop to select the glia in the images and saved the selection. We measured the total area of the selection in pixels. For synaptic quantification 30 synapses per genotype were analyzed.

### Acute brain slice preparations

All procedures for brain slice preparation were performed according to guidelines approved by the Duke University Institutional Animal Care and Use Committee. Briefly, acute brain slices were prepared from mice of both sexes (P18-P25) by deeply anesthetizing via isoflurane inhalation, acute decapitation, and rapidly removing brains for chilling in ice-cold cutting solution. A potassium gluconate cutting solution (pH 7.30) was used for slicing (130 mM K-gluconate, 15 mM KCl, 0.05 mM EGTA, 20 mM HEPES, 25 mM glucose). Acute sagittal slices (250 um) of the cerebellum containing intact Purkinje cells were obtained using a vibrating tissue slicer (VT1200S; Leica) and incubated in ACSF (125 mM NaCl, 26 mM NaHCO3, 1.25 mM NaH2PO4, 2.5 mM KCl, 25 mM glucose, 1 mM MgCl2, 2 mM CaCl2) at 32°C for 20 minutes. Slices were kept at room temperature in ACSF, continuously oxygenated with 95% O2 and 5% CO2. Slice recording conditions: Slices were continuously superfused (3 mL/sec flow rate) with oxygenated ACSF at room temperature. Neurons were visualized using an upright microscope (S-SCOPE-II; Scientifica) equipped with infrared differential interference contrast optics.

### Brain slice electrophysiology

Voltage-clamp recordings were sampled at 50 kHz and bandpass filtered at 1 Hz and 15 kHz. Signals were collected and amplified by a Multiclamp 700B amplifier (Molecular Devices) and digitized using a Digidata 1440A (Molecular Devices). Data was acquired using pClamp 10 (Molecular Devices). A cesium internal solution (pH 7.30) was used for whole-cell recordings (150 mM Cs-gluconate, 5 mM HEPES, 1.1 mM EGTA, 7.5 Phosphocreatine-tris2, 2.5 mM Phosphocreatine-Na2, 3 mM MgATP, 0.5 mM NaGTP, 1.5 mM QX314-Cl, 2 mM TEA-Cl). Pipettes were pulled from borosilicate capillary glass (Warner Instruments) with target tip resistances of 3-9 megaohms when filled with internal solution. Whole-cell, voltage-clamped recordings were performed with membrane holding potentials of -70 mV to isolate EPSCs and 0 mV to isolate IPSCs. For quality control, EPSCs were isolated with bath application of 5 uM gabazine in a subset of recordings to ensure that cells were at the correct holding potentials. Only EPSCs recorded in the absence of gabazine were analyzed to reflect consistent treatment across all cells. Recordings were performed blinded to genotype. For recordings of synaptic activity (Fig. 8 A-H, J-M): WT: 10 animals (5 male, 5 female), KO: 13 animals (8 male, 5 female). For extracellular recordings (Fig. 8 I): WT: 18 animals (7 male, 11 female), KO: 22 animals (14 male, 8 female). Every data point in the summary graphs represents an individual cell recording, with multiple cells from each animal. Electrophysiological analyses were performed using custom scripts in MATLAB (MathWorks).

### Resource Availability

Further information and requests for resources and reagents should be directed to and will be fulfilled by the lead contact, Mary E Hatten. All unique/stable reagents generated in this study are available from the lead contact with a completed materials transfer agreement. There are restrictions to the availability of mice used in this study due to the lack of an external centralized repository for its distribution and our need to maintain the stock. We are glad to share mice with a completed materials transfer agreement and reasonable compensation by request or for its processing and shipping.

## Data Availability

RNA sequencing data has been submitted to GEO under accession number GSE254224. Proteomic datasets will be submitted to ProteomeXchange.

## Notes

### Competing Interest Statement

The authors have declared no competing interest.

