## Supplementary Material for "Mice lacking *Astn2* have ASD-like behaviors and altered cerebellar circuit properties"

### Hanzel et al Supplementary Materials

#### SUPPLEMENTARY FIGURES

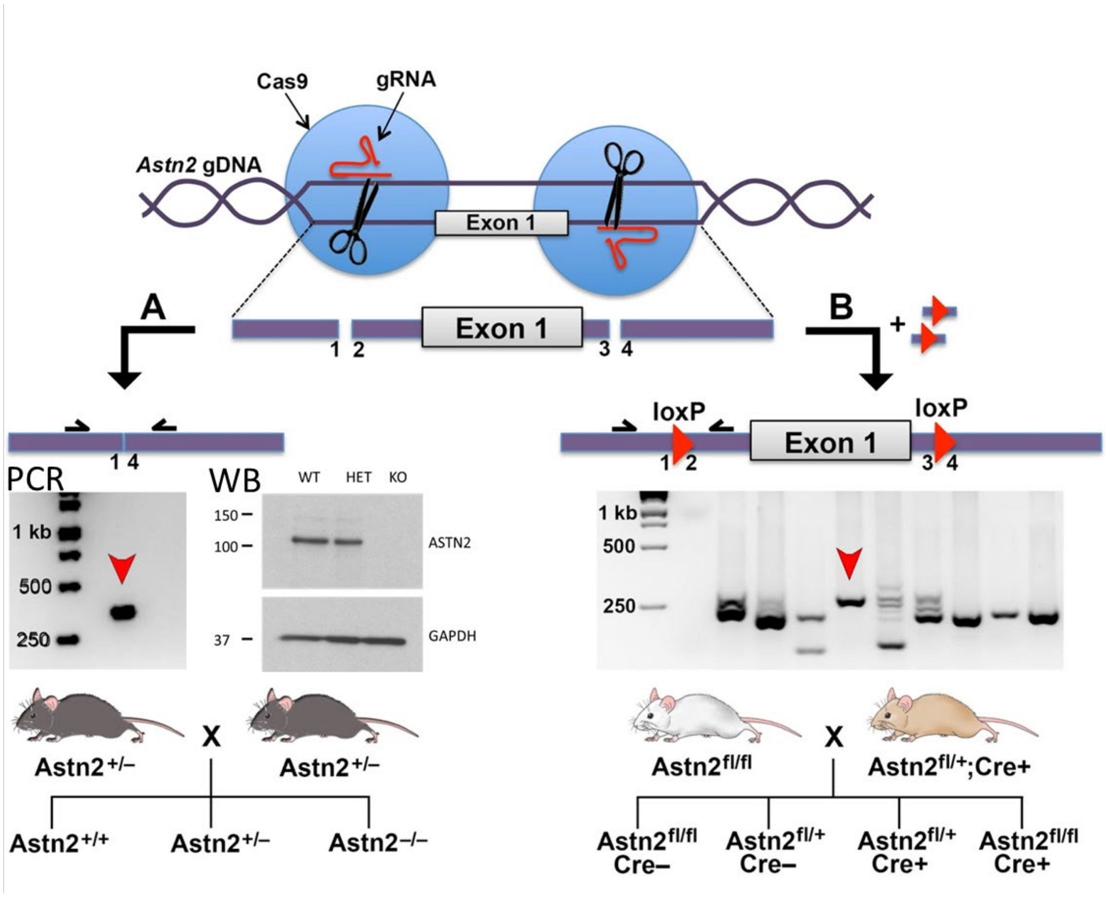


**Supplementary Figure 1: Generation of *Astn2* knockout and conditional knockout mouse.** **A)** The *Astn2* knockout (KO) mouse line was generated using CRISPR-Cas9 technology by targeting the first exon and the promoter region of the *Astn2* gene. To knock out *Astn2*, two guide RNAs flanking the promoter and the exon containing ATG start codon were selected. RNP containing guide RNA and cas9 protein was co-injected into C57BL/6 fertilized embryos. To assess the deletion of *Astn2*, genomic DNA from founders was amplified and sequenced. The loss of ASTN2 protein expression was confirmed using Western blot analysis (WB). **B)** To generate a conditional *Astn2* knockout mouse line, two sgRNAs targeting the exon containing ATG start codon were designed and made. A donor plasmid containing the exon flanked by two loxP sites and two homology arms of 800 bp each was designed and constructed. Genotyping with primers flanking the two loxP sites for potential founder mice were performed. PCR products were analyzed by Sanger sequencing to validate the 5′ and 3′ loxP insertions in the mouse genome.

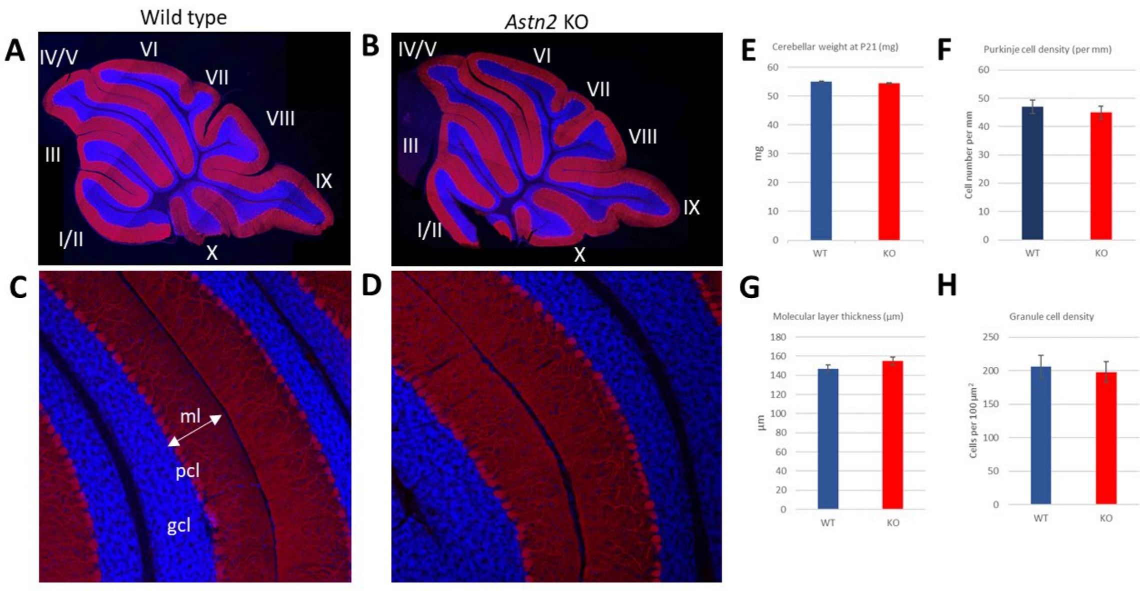


**Supplementary figure 2: *Astn2* KO mice do not have defects in cerebellar gross morphology. A-B)** Midsagittal sections through P21 (A) wild type and (B) *Astn2* KO cerebellum stained with antibodies against calbindin (Purkinje cells) and Hoechst (cell bodies). *Astn2* KO cerebellum has a normal shape and size and contains all expected lobules. **C-D)** 20x image of cerebellar layers in (C) WT and (D) *Astn2* KO animals. Measurements of layer thickness and cell numbers were performed at P21. **E)** Cerebellar weight at P21 is comparable between genotypes. **F)** Purkinje cell density is comparable between genotypes. **G)** Molecular layer thickness is comparable between genotypes. **H)** Granule cell density in the granule cell layer is comparable between genotypes. Data is presented as the mean + SEM. Data analyzed with Student's T test $*p < 0.05$

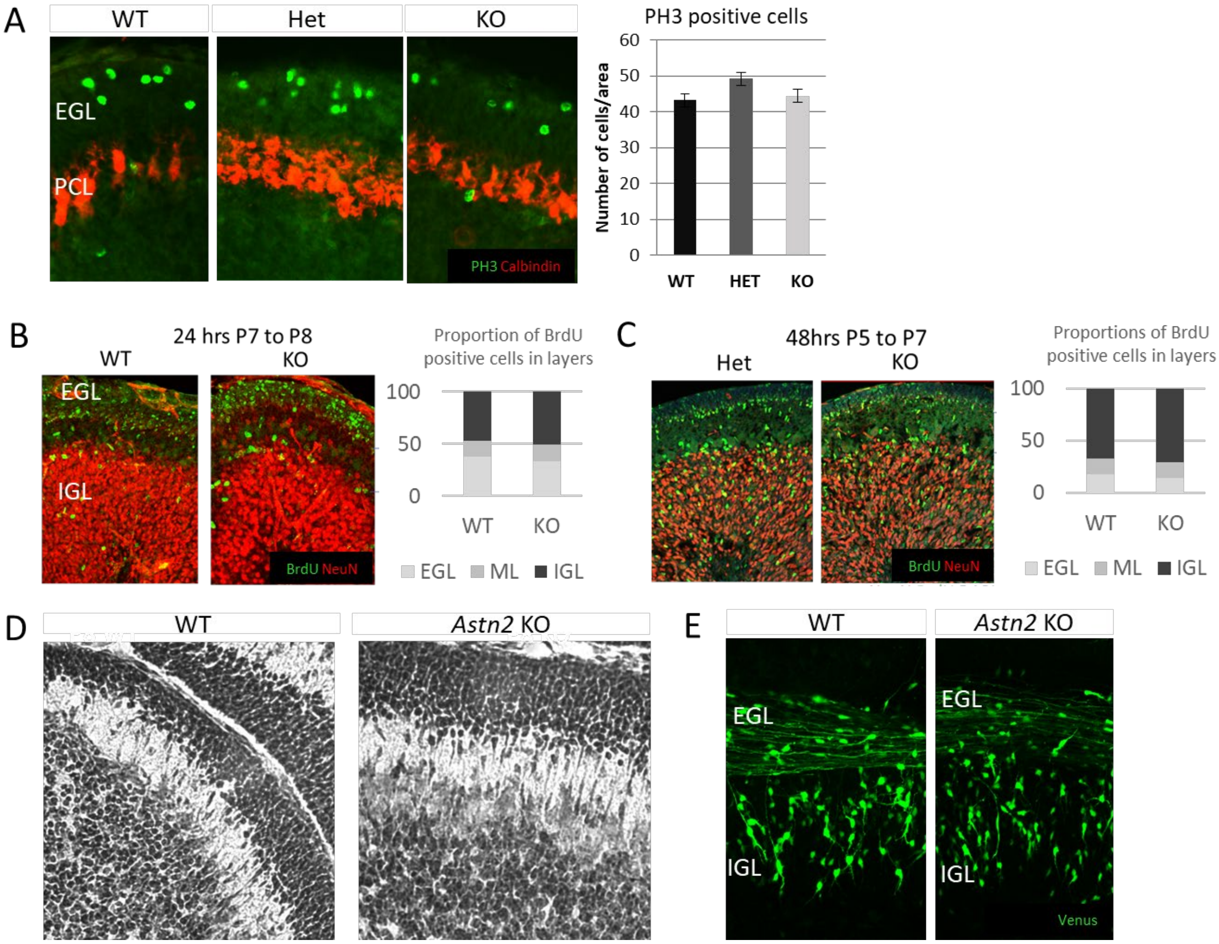


**Supplementary figure 3: No abnormalities in cerebellar granule cell proliferation or migration in *Astn2* KO mice. A)** P7 wild type, heterozygous and *Astn2* KO cerebella were immunostained for PH3 and calbindin. Immunostaining for PH3, a mitosis marker, reveals no differences between genotypes in proliferation rate of granule cell precursors. **B)** P7 animals were injected with BrdU and sacrificed 24hr later to measure the rate of migration of newly differentiated granule cells. The proportions of BrdU positive cells in cerebellar layers (EGL = external granular layer; ML = molecular layer; IGL = internal granular layer) were comparable between genotypes. **C)** BrdU injection at P5 and sacrifice after 48 hrs also did not reveal differences in granule cell proliferation and migration dynamics. **D)** Cresyl violet staining did not reveal any differences in the EGL thickness, the numbers or the shape of the migrating granule cells in the molecular layer or numbers of granule cells in the IGL. **E)** Electroporation of P7 cerebellum with Venus did not reveal any differences in the numbers or morphology of migrating granule cells in cerebellar slices 48 hours post electroporation.

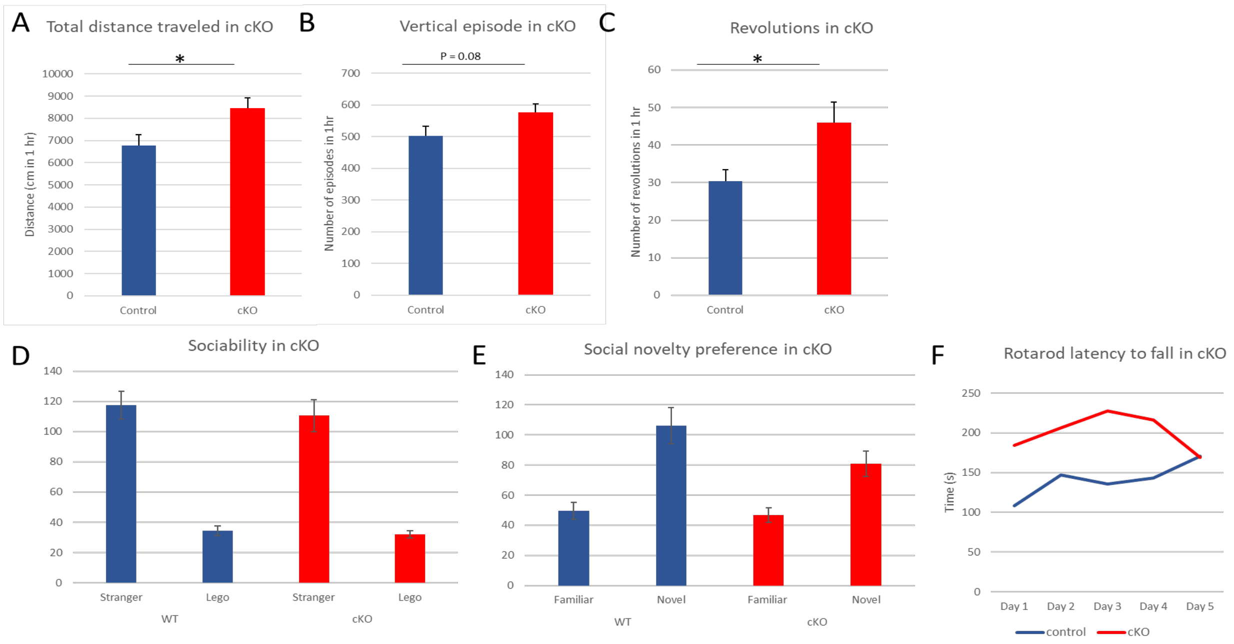


**Supplementary figure 4: Purkinje cell-specific *Astn2* conditional KO animals show hyperactivity and some repetitive behaviors in the open field but no social or motor coordination deficits. A-C)** Open field experiments were performed on 8-10 week old Pcp2-cre *Astn2$^{flox/flox}$* animals (n = 13) and control littermates (n = 14). (A) The total distance traveled was increased significantly in the *Astn2* cKO animals (p = 0.023). (B) There was a trend towards an increase in the number of vertical episodes (rearing) in the *Astn2* cKOs as compared to WTs and hets (p = 0.08). (C) The number of revolutions (circling) was significantly increased in the *Astn2* KOs (p = 0.025). **D-E)** The three chamber social test was performed on 8-12 week old *Pcp2-cre Astn2$^{flox/flox}$* animals (n = 12) and control littermates (n = 9). *Astn2* cKO animals show a preference for the social stimulus similar to control littermates. *Astn2* cKO animals interact with the novel animals significantly longer than with the familiar animals similar to controls. **F)** 8-12 week old control (n = 4) and *Pcp2-cre Astn2$^{flox/flox}$* (n = 4) animals were placed on an accelerating rotarod for five consecutive days. An average of three trials per day was recorded. Time to fall from the rotarod was measured. *Astn2* cKO animals did not show significant decrease in their latency to fall compared to control animals. Data is presented as the mean + SEM. Data analyzed with Student's T test; *p < 0.05

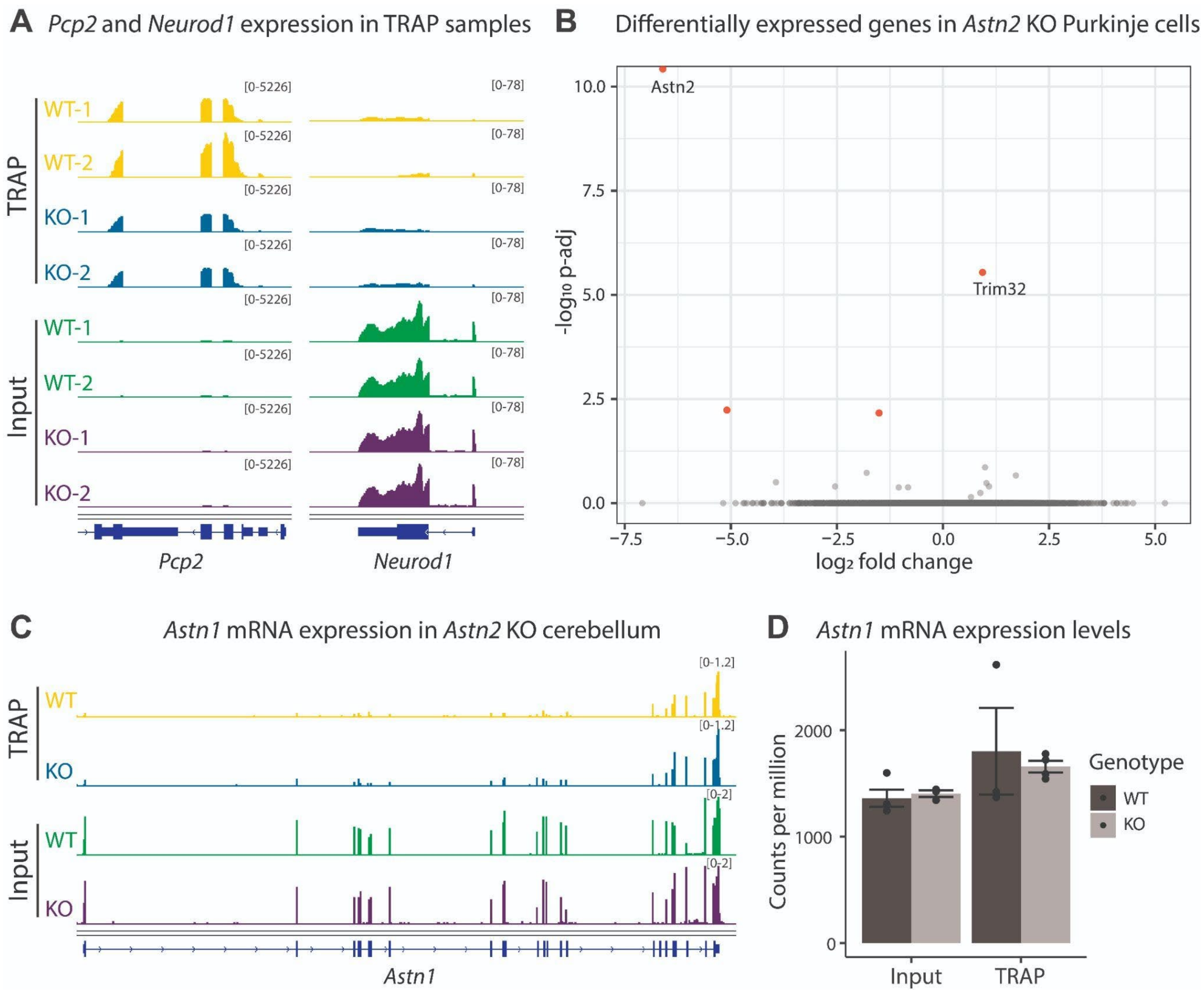


**Supplementary Figure 5. Gene expression changes in *Astn2* KO Purkinje cells. A)** Genome browser view of representative RNA-seq tracks from WT and *Astn2* KO TRAP and input samples. Purkinje cell marker Pcp2 is enriched in TRAP samples, while granule cell marker Neurod1 is enriched in input samples. **B)** Volcano plot depicting differentially expressed genes (*P*-adj<0.05, indicated with red) in *Astn2* KO Purkinje cells, compared with WT littermates, identified using DESeq2. N=3 WT and 4 KO. **C)** Genome browser view of *Astn1* gene from representative WT and *Astn2* KO samples. **D)** *Astn1* mRNA expression levels in WT and *Astn2* KO samples (counts per million). Error bars show mean ± sem. N=3-4/group.

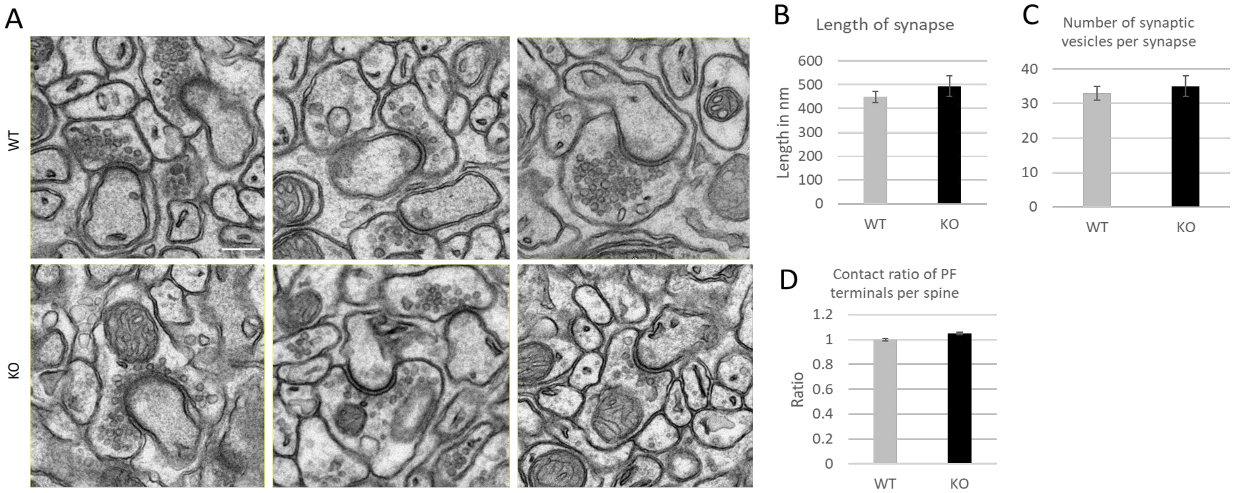


**Supplementary figure 6: No differences are found in the ultrastructure of the synapse in the *Astn2* KO animals. A)** Example electron microscopy images showing synapses between granule cells and Purkinje cells in the molecular layer of wild type (*top panel*) and *Astn2* KO (*bottom panel*) cerebellum at P22. **B)** The length of the synapse measured as the contract between the pre and post synaptic compartments was not changed in *Astn2* KOs. **C)** The number of synaptic vesicles in the synapse was not changes in *Astn2* KO cerebellum. **D)** The contract ratio of PF terminals per PC spine was not changes between genotypes. Data is presented as the mean + SEM. Data analyzed with Student's T test; $*p < 0.05$

### SUPPLEMENTARY METHODS

*BrdU assay*

P7 control and *Astn2* KO mice were injected with 50 μg of BrdU in PBS per gram of body weight in the back of the head over the cerebellum, sacrificed one hour later, and the cerebella were removed and fixed in 4% PFA/PBS overnight. The cerebella were then washed in PBS, embedded in 3% agarose in PBS, and 50 μm sagittal sections were made using a Leica VT1000S vibratome set at a speed of 3 and frequency of 6. The sections were processed for BrdU staining using a BrdU Labeling and Detection Kit II (Roche) and stained with calbindin to label Purkinje cells. The sections were then imaged using confocal microscopy. The number of BrdU positive cells and total GCPs in the EGL were counted and the percentage of BrdU positive cells was calculated.

*Electroporation of Constructs into Cerebellum and Organotypic Slices*

Venus cDNA was PCR generated with the following primers: Venus 5' EcoRI primer GAGAAGGAATTCACC ATGGTGAGCAAGGGCGAGGAG and Venus 3' NotI primer – stop GAGAAGGCGGCCGC TTACTTGTACAGCTCGTCCATGCCG. The cerebella were dissected out at P7 in HBSS containing 2.5 mM HEPES (pH 7.4), 46 mM D-glucose, 1 mM CaCl2, 1 mM MgSO4, 4 mM NaHCO3, and Phenol Red (hereto referred to as HBSS with extra glucose) on ice. The dissection medium was then removed and DNA was diluted to 0.5 μg/μl in HBSS with extra glucose. The PCR product was inserted into pCIG2 (provided by Dr. Franck Polleux) digested with EcoRI and NotI. The cerebella were soaked in the DNA for 15-20 minutes on ice, and were then transferred one at a time into the well of an electroporation chamber (Protech International Inc. CUY520P5 platinum electrode L8xW5xH3 5mm gap) that was placed on ice. The cerebella were electroporated dorsal to ventral for 50 ms at 80 V, for a total of 5 pulses with an interval of 500 ms between pulses, using an electro-square-porator, ECM 830 (BTX Genetronics). The cerebella were then removed from the chamber and placed on ice to recover for 10 Minutes. Subsequent to electroporation the cerebella were embedded in 3% agarose in HBSS containing 30 mM D-glucose, and 250 μm coronal slices were made using a Leica VT1000S vibratome set at a speed of 3 and frequency of 6. Slices were then placed on MILLICELL CM 0.4 um culture plate inserts in a 6 well plate with 1.5 ml of culture medium (BME, 25 mM D-glucose/1x Glutamine/1x ITS/1x Pen-Strep) below the insert. The organotypic slices were incubated at 35°C/5% CO2 for 48 hours before fixing with 4% PFA/4% sucrose/PBS for 2 hours at room temperature. Slices were mounted on slides and imaged by confocal microscopy.
